# Exercise Training Remodels Inguinal White Adipose Tissue Through Adaptations in Innervation, Vascularization and the Extracellular Matrix

**DOI:** 10.1101/2022.08.09.503375

**Authors:** Pasquale Nigro, Maria Vamvini, Jiekun Yang, Tiziana Caputo, Li-Lun Ho, Danae Papadopoulos, Nicholas P. Carbone, Royce Conlin, Jie He, Michael F. Hirshman, Joseph D. White, Jacques Robidoux, Robert C. Hickner, Søren Nielsen, Bente K. Pedersen, Manolis Kellis, Roeland J. W. Middelbeek, Laurie J. Goodyear

**Affiliations:** Section on Integrative Physiology and Metabolism, Joslin Diabetes Center, Harvard Medical School, Boston, MA; Division of Endocrinology, Diabetes and Metabolism, Beth Israel Deaconess Medical Center, Harvard Medical School, Boston, MA; Computational Science and Artificial Intelligence Laboratory, Massachusetts Institute of Technology, Cambridge, MA; Department of Pharmacology and Toxicology, East Carolina University, Greenville, NC; Department of Nutrition and Integrative Physiology, Florida State University, Tallahassee, FL; The Centre of Inflammation and Metabolism and the Centre for Physical Activity Research, Rigshospitalet, University of Copenhagen, Denmark

**Keywords:** Exercise, White adipose tissue, ECM, Innervation, Vascularization, NEGR1, PRDM16, Spatial transcriptomics, Proteomics, Adipo-Clear

## Abstract

Inguinal white adipose tissue (iWAT) is essential for the beneficial effects of exercise training on metabolic health. Extracellular matrix (ECM) composition, innervation, and vascularization are all important regulators of iWAT function, yet whether exercise training improves these structural components of iWAT is unknown. Using biochemical, imaging, and multi-omics analyses we find that 11-days of wheel running in male mice causes profound iWAT remodeling including decreased ECM deposition and increased vascularization and innervation. We identify adipose stem cells as the main contributors to training-induced ECM remodeling, determine that training causes a shift from hypertrophic to insulin-sensitive adipocyte subpopulations, show that the PRDM16 transcriptional complex is necessary for iWAT remodeling and beiging, and discover neuronal growth regulator 1 (NEGR1) as a link between PRDM16 and neuritogenesis. Exercise training leads to remarkable adaptations to iWAT structure and cell-type composition that can confer beneficial changes in tissue metabolism.

**Graphical Abstract:** 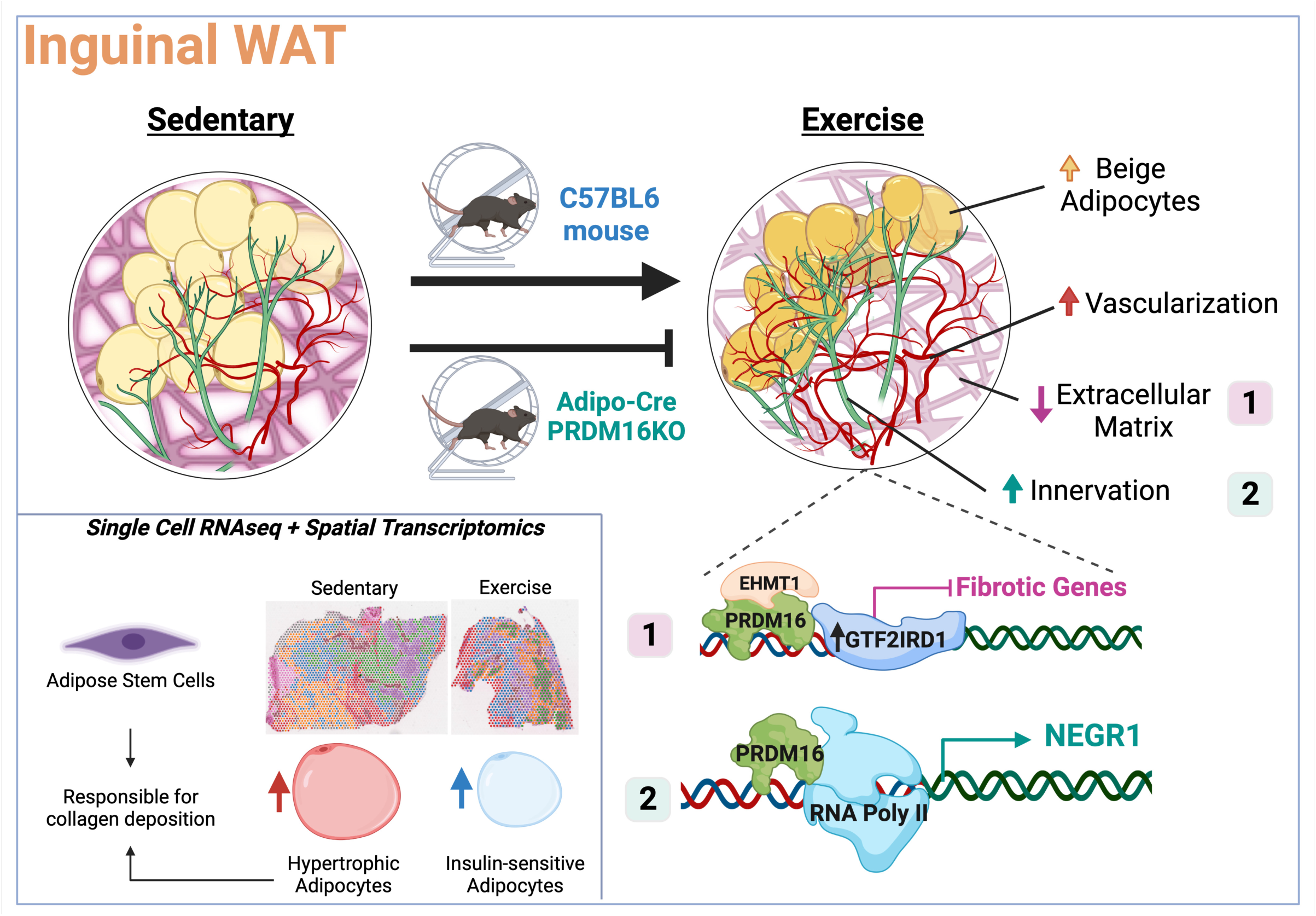

## INTRODUCTION

White adipose tissue (WAT) is a heterogeneous and highly dynamic endocrine organ that is responsive to numerous stimuli and is capable of undergoing remodeling to meet the metabolic and energy demands of the body (Li et al., 2021). In obesity, homeostasis of both subcutaneous and visceral WAT depots is impaired, contributing to systemic metabolic disease (Ruiz-Ojeda et al., 2019). Pathological expansion of the extracellular matrix (ECM), excessive tissue fibrosis, inflammation and dysregulated lipid accumulation have been established as important factors in this dysregulation of WAT (Ruiz-Ojeda et al., 2019). In contrast to obesity, exercise training can improve systemic metabolic homeostasis, and recent data suggest that some of these beneficial effects of exercise are mediated through adaptations to the subcutaneous inguinal WAT (iWAT) (Yang et al., 2022; Nigro et al., 2021; Takahashi et al., 2019; Stanford et al., 2015; Trevellin et al., 2014; Lira et al., 2014; Gollisch et al., 2009;). The underlying molecular and cellular mechanisms mediating the effects of exercise training on iWAT are still poorly understood.

ECM is a highly dynamic structure; continuously modified, degraded and deposited (Marcelin and Clément, 2021). ECM remodeling affects important cellular functions such as adhesion, migration, proliferation, and differentiation, and is also closely linked with fundamental tissue processes including vascularization (Lu et al., 2011) and innervation (Long and Huttner, 2019). Dysregulated ECM remodeling is found in numerous conditions including inflammatory diseases, heart disease, and cancer, and given this clinical relevance, research aimed at understanding the cellular mechanisms of pathological ECM remodeling has become more prominent. ECM remodeling in WAT has been studied mainly in the context of obesity. In WAT, ECM turnover is associated with the formation of new blood vessels that ensures the healthy expansion of the tissue by preventing tissue hypoxia, fibrosis and WAT dysfunction (Spencer et al., 2011). There is also a pivotal role of sympathetic innervation in the regulation of WAT physiology, remodeling and adaptations (Lambert et al., 2015). For example, stimulation of sympathetic innervation in WAT following cold exposure is required for beige cell formation and non-shivering thermogenesis (Bartness et al., 2014; Cao et al., 2019), both critical in maintaining systemic glucose homeostasis. Thus, understanding ECM remodeling, and the associated cellular processes of angiogenesis and innervation may facilitate the development of new therapeutic targets for numerous diseases.

In contrast to the investigation of ECM with obesity, the effect of exercise training on ECM remodeling in iWAT has not been previously studied, and so it is not known if ECM is essential for the beneficial effects of training on this tissue. Similar to ECM remodeling, there are also no reports investigating the effects of exercise training on iWAT innervation. Exercise training has been shown to increase the expression of the angiogenic factor *Vegfa* in scWAT (Lee, 2018; Stanford et al., 2015), suggesting that training promotes iWAT vascularization, although this has not been directly investigated. Thus, while exercise training is known to result in a more favorable metabolic profile in iWAT, it is essential to determine if structural components of iWAT adapt in a beneficial manner to exercise training, as these could provide the basis for novel therapeutic approaches.

Here, we test the hypothesis that exercise training alters ECM deposition and induces innervation and vascularization in iWAT, and determine the molecular and cell type specific adaptations to iWAT in response to exercise. Using a combination of biochemical, imaging, genetic, and multi-omics approaches including advanced spatial transcriptomics, we find that exercise training dramatically reduces ECM stiffness, increases vascularization, and promotes sympathetic innervation with neuronal refinement in iWAT. Using single cell transcriptomics, we identify the predominant cell populations that contribute to iWAT ECM physiology and determine how exercise affects collagen deposition by modulating both the cell composition and gene expression profile within specific cell types in iWAT. We also determine that PRDM16 is a regulator of training-induced beiging in iWAT and identify Neural Growth Regulator 1 (NEGR1) as a link between PRDM16 and exercise-induced neuritogenesis. Thus, exercise training is a potent physiological stimulus for beneficial remodeling of iWAT.

## RESULTS

### Exercise training reduces collagen deposition in iWAT

To determine the effects of exercise training on the iWAT ECM, 8-week old male C57BL/6 mice were fed a standard diet (20% kCal from fat) and housed in individual cages with (exercise training) or without (sedentary) voluntary access to a running wheel. Exercise trained mice ran 6.0±1.6 km/day and had increased food intake compared to sedentary mice (Figure S1A). Sedentary mice increased their body weight by 7% during the 11 days whereas exercise trained mice maintained their body weight, although there was no significant difference between the two groups at day 11 (Figure S1B). Exercise training resulted in improved fasting blood glucose, insulin, and HOMA-IR (Figures S1C-E). At the tissue level, exercise training reduced fat mass (Figure 1A). The difference in appearance and size of the trained fat pad was apparent under 4x magnification (Figure 1B, left panels), and abundance of smaller and multilocular adipocytes in trained iWAT was observed using 20x magnification (Figure 1B, right panels). To evaluate tissue remodeling, we measured changes in collagen, the main component of ECM. We found decreased collagen content in trained iWAT compared to sedentary iWAT using a hydroxyproline assay (Figure 1C). This finding was corroborated by microscopic imaging using Sirius Red staining for collagen where trained iWAT showed thinner and fewer collagen structures (Figure 1D). Using the TWOMBLI software package (Wershof et al., 2021), we determined that trained iWAT showed a decreased collagen area (Figure 1E), reduced ‘high density matrix’(HDM) (Figure 1F), and a reduction in the total length of collagen branches compared to sedentary iWAT (Figure 1G). These findings support the hypothesis that exercise training causes ECM remodeling through a decrease in iWAT collagen deposition and ECM stiffness.

**Figure 1.**
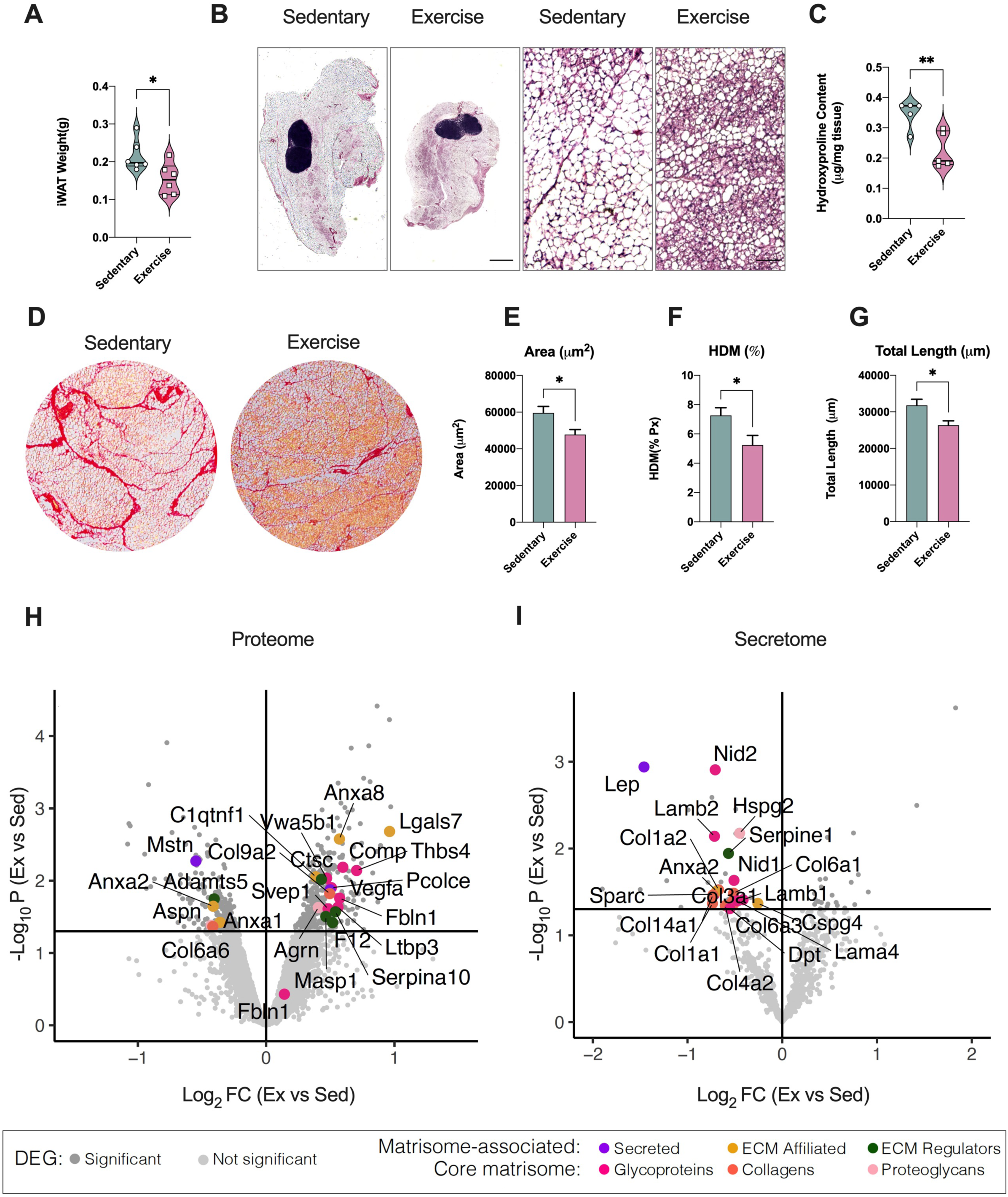
Exercise training reduced ECM deposition in iWAT, modulating Matrisome-associated and Core Matrisome gene expression levels. (A) Inguinal subcutaneous white adipose tissue (iWAT) mass of male sedentary and trained mice (*n=6/group*). (B) Representative H&E staining images of iWAT from sedentary and trained mice. Scale bar: 1000 µm (4x) and 50 µm (20x), respectively. (C) Hydroxyproline content in iWAT from sedentary and trained mice (*n=5/group*). (D)Representative Sirius Red staining images of iWAT from sedentary and trained mice. Scale bar: 100 µm (10x). (E-G) Quantification of ECM deposition pattern area (E), percentage of high-density matrix (HDM) (F) and total length of fibers (G) in iWAT (*n=4/group; calculated from 8 fields/mouse*). (H)Volcano plot displaying up- and down-regulated proteins with exercise training detected in iWAT (iWAT proteome analysis). Lines are drawn to define restriction of log_2_ FC value of 0.5 and −log_10_ of p-value 0.01. Significant proteins are colored based on matrisome subcategories. (I) Volcano plot displaying up- and down-regulated proteins with exercise training detected in iWAT conditioned media (secretome analysis). Lines are drawn to define restriction of log_2_ FC value of 0.5 and −log_10_ of p-value 0.01. Significantly proteins are colored based on matrisome subcategories. Data are presented as mean ± SEM and were compared using unpaired two-tailed Student’s t-test. *p < 0.05, and **p < 0.01.

### Identification of exercise-modulated ECM genes expressed in iWAT

To determine gene expression changes driving exercise-induced iWAT ECM remodeling, we utilized our previously published microarray dataset generated using iWAT from sedentary mice and 11-day-wheel-trained mice (GSE68161) (Stanford et al., 2015). We performed pathway enrichment analysis using the murine ECM proteome or ‘matrisome’ database (Naba et al., 2012, 2016) which includes not only all the genes encoding structural ECM components (‘Core matrisome’), but also the genes encoding proteins that can interact with and remodel ECM (‘Matrisome-associated’). For Matrisome-associated genes, 689 out of 837 or 82% were expressed in iWAT, and for Core matrisome 234 out of 274 or 85% of genes were expressed in iWAT (Figure S1F). Of these genes expressed in iWAT, we found that 16% of Matrisome-associated and 18% of Core Matrisome genes were either significantly up-regulated or down-regulated by exercise training (Figure S1F and Table S1), underscoring that there are robust exercise-induced transcriptional changes of ECM genes.

Matrisome-associated proteins contain ECM regulators, ECM-affiliated proteins, and secreted factors that may interact with core ECM proteins. Figure S1G summarizes the top 10 genes with the greatest absolute fold change for each of these three categories. Exercise training decreased ECM regulators that are known for their inhibitory role in WAT ECM remodeling (*Timp4*) (Maquoi et al., 2002) or their negative effect on the beiging process (*Adamts5*) (Bauters et al., 2017; Bondeson et al., 2008). For ECM affiliated genes, exercise training decreased genes implicated in the development of WAT fibrosis (*Galectin-3*, *Lgals3*), (Martínez-Martínez et al., 2016) and increased genes involved in neuronal network regulation and neuron myelination (*Neuron glial antigen 2; NG2/CSPG4*), (Sakry et al., 2014; Sakry and Trotter, 2016). A subset of Matrisome-associated genes is annotated as secreted factors. Of these, exercise training down-regulated Leptin (*Lep*), a marker of adipose tissue mass, and *S100a8,* a potent mediator of fibrosis and inflammatory response (Tammaro et al., 2018), and increased genes involved in the beiging process and in adipose tissue innervation (*Nrg4* and *S100b*)(Wang et al., 2014; Zeng et al., 2019).

The core matrisome is composed of collagens, ECM glycoproteins, and proteoglycans. Among the genes included in the iWAT core matrisome, 29 genes were significantly down-regulated, and 13 genes were significantly up-regulated by exercise training (Figure S1F and Table S1). Matrisome enrichment analysis revealed that out of the 29 down-regulated genes 5 are collagens, 18 are glycoproteins, and 6 are proteoglycans (Figure S1H). To map the exercise-regulated core matrisome genes to the different compartments of ECM, we generated an iWAT ECM protein-protein interaction (PPI) network in Cytoscape integrating our data with publicly available data (Tsutsui et al., 2021) (Figure S1I). The collagen core components down-regulated by exercise training (*Col4a1*, *Col4a2*, *Col18a1*, *Col23a1*) are mainly located in the basement membrane compartment while the proteoglycans (*Fmod*, *Bgn*,*Vcan*, *Srgn*) and glycoproteins (*Mfge8*, *Mfap3*, *Mfap5*, *Efemp1*, *Fbn1*, *Fbln2*) are expressed in the interstitial matrix. We validated the effects of exercise training on mRNA expression of selected genes from matrisome-associated and core matrisome using qRT-PCR (Figure S1J-K). Taken together, these results suggest that exercise training induces ECM remodeling through modulation of expression of matrisome-associated and core matrisome genes, which are also known to regulate other tissue processes such as innervation and beiging.

### Quantitative proteomic profiling revealed iWAT ECM remodeling with exercise training

The total proteome of iWAT from sedentary and trained mice was quantified using high resolution mass spectrometry. We detected a total of 6,910 proteins in iWAT. Of these proteins, 90% were predicted to have an intracellular localization, whereas ~10% were annotated as extracellular proteins. A total of 482 (7%) proteins were differentially expressed between sedentary and trained mice (p<0.05, Table S2), with training up-regulating 300 proteins and down-regulating 182 proteins. Of these, 23 proteins were annotated for matrisome, 17 up-regulated and 6 down-regulated (Figure 1H).

Gene Ontology (GO)-Cellular Component (GO-CC) enrichment analysis of the iWAT showed that training led to proteomic changes in all major cellular components (Figure S1L). Mitochondria was the cellular component most affected by exercise training with 10% proteins increased and 4% proteins decreased (Figure S1L). We mapped the exercise-regulated proteins to the murine ECM proteome or ‘matrisome’ database (Naba et al., 2012, 2016). This analysis showed that exercise increased the matrisome-associated pathways and decreased the core matrisome and collagen-associated pathways (Figure S1M).

While tissue proteomics can identify annotated extracellular proteins, it does not provide confirmation of protein release from iWAT. Thus, to determine secreted factors involved in iWAT ECM remodeling we performed a conditioned media secretome analysis of iWAT from sedentary and trained mice. We identified 726 proteins of which 8% (n=60) were differentially expressed with exercise training (Figure 1I, Table S3). Exercise training up-regulated 23 and down-regulated 37 proteins (Figure 1I). Fifty percent (n=19) of the down-regulated proteins were ECM related genes and no ECM genes were significantly up-regulated in the secretome analysis. GO-CC enrichment analysis revealed that exercise training down-regulated a large number of proteins localized in the extracellular space (Figure S1N). Further pathway analysis of the extracellular down-regulated proteins showed enrichment for the ECM core matrisome (Figure S1O). The most significantly down-regulated core matrisome proteins included leptin (*LEP*), annexin A2 (*ANXA2*), the glycoprotein *SPARC*, and the collagen *COL1A1* (Figure 1I). By mapping these proteins using PPI network analysis, we determined that most of the down-regulated matrisome proteins are located in the basement membrane layer of ECM (Figure S1P). Overall the proteome and secretome analyses of iWAT corroborate the transcriptomic findings that exercise training down-regulates protein level and secretion of collagen species, especially the proteins found in the basement membrane.

### Spatial and single-cell transcriptomics identifies main contributors of ECM in iWAT

To identify which adipose tissue cell types are mainly responsible for ECM remodeling, we re-analyzed previously published single-cell RNA sequencing datasets of both adipose stromal-vascular and mature adipocyte fractions (Yang et al., 2022; Rajbhandari et al., 2019). The cell populations include mature adipocytes (both white and beige), preadipocytes, adipose stem cells (ASCs), lymphoid and myeloid immune cells, vascular cells, smooth muscle cells, glial cells and epithelial cells (Figure S2A and S2C). In these data, the down-regulated ECM secreted proteins that we detected in the multi-omic analysis (Figure 1) were mostly expressed in ASCs, preadipocytes, mature adipocytes, glial cells, and vascular cells (endothelial, smooth muscle, lymphatic endothelial cells and pericytes) (Figure S2B, S2D). Since the ASC/preadipocytes cluster showed the greatest consistency in terms of the expression of ECM proteins across the single-cell RNA datasets, we estimated transcription factor (TF) activity in these cells by averaging gene expression of curated targets for each TF. ASCs demonstrated the highest TF activity compared to other cell types with SMAD3 being one of the TFs exhibiting significantly high TF activity within the ASCs (Figure S2E). Our next goal was to determine the spatial distribution of the cells contributing to ECM. This is especially important because ECM regulates other spatially confined processes such as innervation and beiging. Thus, to determine localization of the cell types in iWAT we performed spatial transcriptomic analysis (Visium Spatial Gene Expression, 10x Genomics) of iWAT from sedentary and trained mice. Each barcoded spot in Visium is ~50 microns in diameter, capturing between 1 and 10 cells/spot based on the detected UMI (unique molecular identifier) (Figure S3A). Exercise trained adipose tissue consists of smaller adipocytes and tissue mass compared to sedentary mice (Figure 1B). Consequently, we observed a smaller number of spots but a larger number of cells/spot in trained mice (Figure S3A). We performed spatial pattern clustering (Figure S3B, C) and scrutinized marker gene expression in the spatially distinct clusters. Based on this, we annotated the major cell types within each spot (Figure 2B). Spots showing a recognizable cell signature aligned with all the cell clusters visualized in the H&E stained images (Figure 2A, B) (e.g. white and beige adipocytes, fibroblast and endothelial/perivascular microglia). Interestingly, exercise training drastically reduced the proportion of fibroblasts, while increasing the proportion of beige, white-beige adipocytes, and endothelial-perivascular microglia cells (Endo/PVM). No differences for ASC and white adipocyte proportions were observed (Figure 2B).

**Figure 2.**
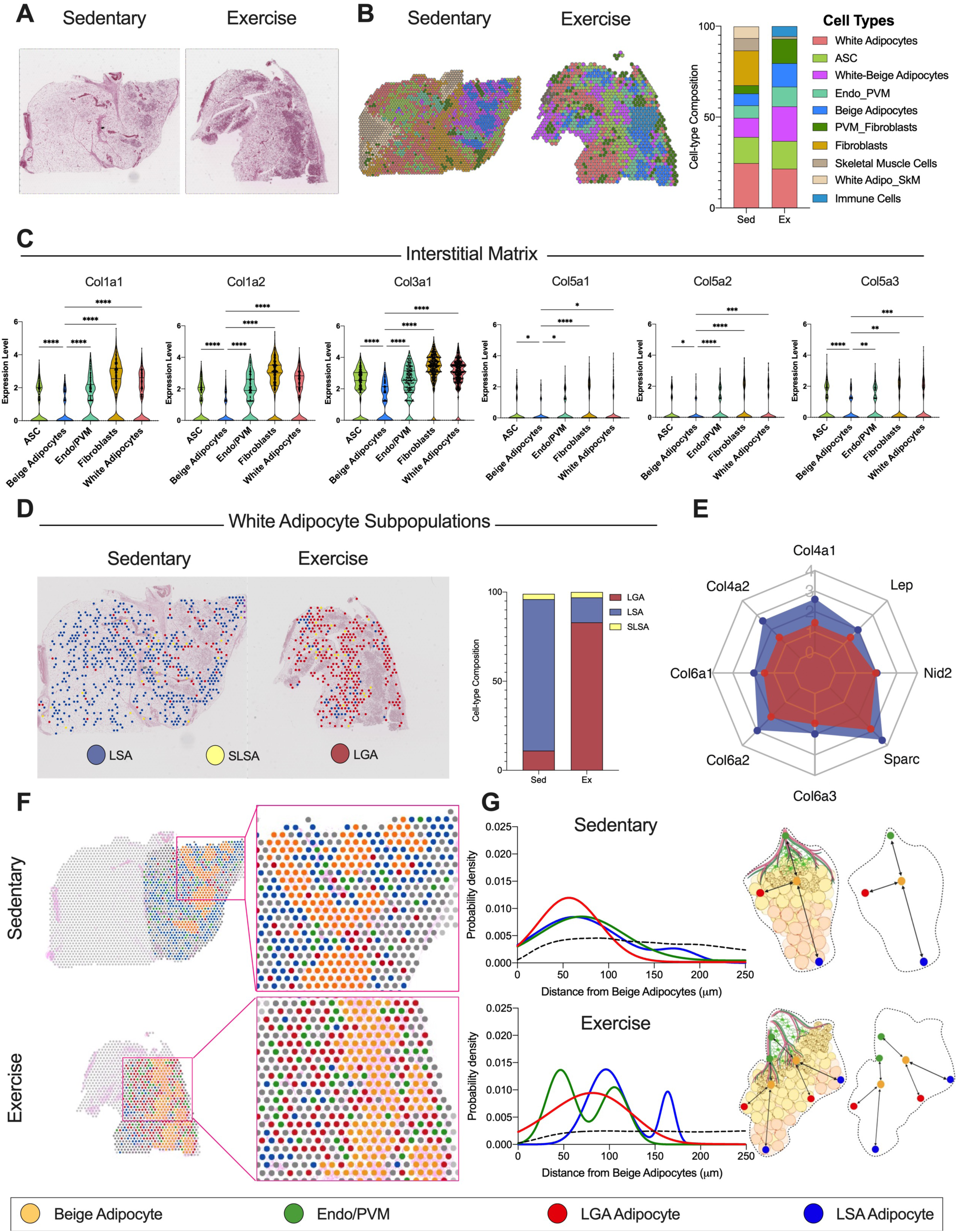
Spatial and single-cell transcriptomics identifies main contributors of ECM in iWAT. (A-B) Magnified view of H&E-stained sections (A) and relative spatial RNA-seq barcoded spot (B) indicating the cell clusters detected in iWAT from sedentary (*left*) and exercise training (*right*) mice. (C) Individual violin plots showing the expression level for interstitial matrix collagens species across 5 selected cell type clusters. (D) Spatial distribution maps showing the three white adipocyte subpopulations lipid-scavenging adipocytes (LSA), stressed lipid-scavenging adipocytes (SLSA) and lipogenic adipocytes (LGA) detected in iWAT from sedentary (*left*) and exercise training (*right*) mice along with the relative proportion plot. (E) Basement membrane components gene expression level in LSA and LGA white adipocyte subpopulations. The gene expression level is presented as a scale ranging from zero to four, where four represents the highest expression level. (F) Spatial distribution maps and relative magnifications showing the selected cell type clusters in iWAT from sedentary (*top*) and exercise training (*bottom*) mice: beige adipocytes (*yellow*), endothelial and perivascular microglia cells (Endo/PVM)(*green*), LGA (*red*), and LSA (*blue*) white adipocyte subpopulations. (G) Probability density distribution plots showing the distance relative to the beige adipocyte for the LGA, LSA and Endo/PVM cluster in iWAT from sedentary (*top*) and exercise training (*bottom*) mice. Dashed black lines indicate the random distribution among the clusters. On the right, a summary cartoon explaining the pattern distribution of the cell clusters. Data are presented as mean ± SEM and were compared using One-way ANOVA. *p < 0.05, **p < 0.01, ***p < 0.001, and ****p < 0.0001.

By comparing expression of different collagen isoforms across the spatially defined clusters, we found that the spots with a cell signature for fibroblasts, ASCs, Endo/PVM and white adipocytes expressed higher levels of *Col1a1*, *Col1a2*, *Col3a1*, *Col5a1, Col5a2,* and *Col5a3* compared to beige adipocytes. All these collagen species are major components of the interstitial matrix ECM compartment. Further, beige adipocytes expressed lower levels of basement membrane components such as *Col6a1*, *Col6a2*, *Nid1*, and *Sparc.* No significant differences were observed for *Col4a1* and *Col4a2* among the 5 spatially defined cell clusters (Figure S3D).

Taken together these data suggest that the mesenchymal stem cells (MSCs) compartment consisting of ASCs, preadipocytes and fibroblasts in iWAT are primarily responsible for collagen synthesis. The reduction of interstitial matrix compartment-related collagens in the trained iWAT was due to a sharp decrease in the proportion of fibroblast and a concomitant large increase in beige adipocytes which exhibit the lowest collagen species content.

### Exercise training promotes a switch in white adipocyte subpopulations

Beside MSCs, mature adipocytes are a major contributor to collagen synthesis and deposition. The expression of collagens in white adipocytes is directly related to their lipid volume; larger cell volume results in a thicker basement membrane ECM compartment (Mariman and Wang, 2010). To test the hypothesis that exercise training directly modulates the expression of collagen species in adipocytes, we annotated the white adipocyte subpopulations using a previously published cell annotation strategy (Sárvári et al., 2021). In iWAT, there are three distinct white adipocyte subpopulations: lipogenic adipocytes (LGA), lipid-scavenging adipocytes (LSA) and stressed lipid-scavenging adipocytes (SLSA). LGA are the insulin sensitive adipocytes involved in *de novo* lipogenesis, while LSA and SLSA are mainly involved in lipid uptake and transport (Sárvári et al., 2021). Interestingly, exercise training shifts the ratio of adipocyte subpopulations from LSA phenotype toward a significant enrichment of LGA (Figure 2D). Few SLSA adipocytes were detected in both sedentary and training iWAT. Across both tissues, LGA showed lower expression of leptin, collagens, and ECM basement membrane components compared to LSA, suggesting that they are the smallest white adipocytes in the tissue (Figure 2E). These findings support the concept that exercise training reduces collagen deposition and synthesis in iWAT.

Our next goal was to understand the cell organization in iWAT and how exercise training impacts spatial organization, an important factor regulating tissue function. We performed a spatial interaction analysis using a previously published pipeline (Shivanandan et al., 2013) to investigate the distance between beige adipocytes and 3 selected cell clusters: Endo/PVM, LGA, and LSA (Figure 2F). For both sedentary and exercise, the number of cells detected and the size of the space within which they were distributed showed a clear pattern with beige cells (Figure 2G,Figure S4B). We also observed that beige adipocytes were closer to LGA than to LSA adipocytes in both conditions. Sedentary iWAT showed all cell clusters to be equally distributed in the selected region of interest. The interaction analysis model provides a strength score that quantifies the probability of cell proximity between beige adipocytes and the other clusters (Probability density). LGA adipocytes had the highest probability density to beige adipocytes with an estimated strength score of 8.02 which reflects nonrandom close proximity (Figure S4B). In contrast, LSA adipocytes were at a greater distance (wide distribution) with an estimated strength score of 5.76 (Figure S4B). Interestingly, trained iWAT showed two peaks for the probability density. For one peak there was a reduced distance between beige adipocytes and the Endo/PMV cluster (strength score=8.26) with training, while the second peak matched the sedentary. The reduced distance peak may be due to the induction of vascularization and innervation in iWAT following training. LSA adipocytes also showed a bimodal distribution, but their location is shifted farther from beige adipocytes (Figure 2G, Figure S4B). Taken together these results indicate that exercise training robustly alters the proportion of LSA to LGA and modulates cell organization, providing an explanation as to how exercise training affects the basement membrane collagen deposition of the white adipocytes.

### Exercise training increases vascularization in iWAT

While we observed changes in cell type interactions with exercise training, we next investigated whether these changes in cell type interactions translated to changes in cellular processes such as innervation and vascularization. According to a gene annotation platform, a significant proportion of the ECM proteins differentially expressed with exercise training in our tissue proteomics and secretome analysis were also annotated in the vascularization and innervation pathways (tissue proteome: 46 out of 166; secretome: 11 out of 23) (Figure S5A,B) (Zhou et al., 2019). These pathways highlight an important role of ECM proteins in neuronal and vascular remodeling in the context of exercise training.

To determine the effects of exercise training on vascularization, angiogenesis of iWAT from sedentary and trained mice was determined using immunofluorescence staining. We used the griffonia simplicifolia lectin (GSA-I) stain which specifically binds to the blood vessel endothelial cells (Laitinen, 1987; Lu et al., 2011) (Figure S5C). Analysis of iWAT images using Cell Profiler (Lamprecht et al., 2007) revealed a significantly higher total capillary density (Figure S5D) and capillary-to-adipocyte ratio (Figure S5E) in the trained iWAT compared with the sedentary iWAT. This exercise-induced iWAT vascularization was also evident at the gene expression level. We found that exercise training induced the expression of pro-angiogenic mediators (Rudnicki et al., 2018), including vascular endothelial growth factor A (*Vegfa*) and its receptor (*Kdr/Vegfr2*), the Notch ligand Jagged-1 (*Jag1*), Angiopoietin-1 (*Angpt1*) and Angiopoietin-2 (*Angpt2*) (Figure S5F-J). These microscopy and gene expression data demonstrate that exercise training increases vascularization in iWAT.

### Exercise training increased sympathetic neurite density in iWAT

The effects of exercise training on iWAT innervation were determined with immunofluorescence analysis using two different stains: synaptophysin (*Syp*), an integral protein on pre-synaptic vesicles and a pan-marker for neural structures, and tyrosine hydroxylase (*Th*) which is a specific marker for sympathetic fibers (Cao et al., 2018). Confocal microscopy revealed that exercise training increased the neural projections in iWAT (Figure 3A) with increased synaptophysin density and synaptophysin/adipocyte ratio (Figure 3B,C). Similarly, staining with TH showed that exercise training increased sympathetic innervation (Figure 3D-F). We performed ‘Adipo-Clear’, which is a whole mount tissue staining that provides a high resolution volumetric fluorescent imaging of the neural network using TH staining (Chi et al., 2018a) and found a tremendous increase of sympathetic innervation with exercise training compared to sedentary iWAT (Figure 3L). In addition, we clearly observed an increase in neurite development, also known as neuritogenesis (Figure 3L ROI). We then measured expression of genes known to be involved in iWAT innervation. Exercise led to higher gene expression levels of the adrenergic receptor beta-2 (*Adrb2*) and beta-3 (*Adrb3*), as well as the purine receptor *P2rx5*, a cell surface marker for beige/brown adipocytes that increases upon β3 stimulation (Ussar et al., 2014) (Figure 3G-J). Cannabinoid 1(*Cnr1*) and the diacylglycerol lipase alpha (*Dagla*), involved in the retrograde endocannabinoid signaling known for its regulatory role in synaptic plasticity, were also found elevated in iWAT following exercise training (Castillo et al., 2012) (Figure 3G-K). At the protein level, we used our iWAT proteomics data to identify the exercise up- and down-regulated proteins involved in the innervation pathways. We performed a PPI network analysis to explore key protein interactions for innervation. Within the up-regulated proteins we identified three significant clusters based on protein-to-protein interaction (cluster 1 to 3) representing the main stages of iWAT neural network adaptations to exercise training (Figure S5K). Proteins in cluster-1 are involved in the *VEGFA*-*EGFR* signaling pathway, which is important for the neurovascular remodeling and neural stem cell activation (Liu et al., 2019). Cluster-2 includes proteins such as paired related homeobox protein 1 (*Prrx1*) *(Shimozaki et al., 2013)* and leukemia inhibitory factor receptor (*Lifr*) (Oshima et al., 2007), which are implicated in the self-renewal of adult murine neural progenitor cells (NPCs). Proteins in cluster-3 are implicated in the clearance of apoptotic neural cells, such as the scavenger receptor class F member 1 (*Scarf1*) and the H-2 class I histocompatibility antigen (*H2-K1*). Similar analysis of the down-regulated proteins revealed a large cluster (cluster-4) (Figure S4L), which includes proteins such as myelin proteolipid protein (*Plp1*) that is a component of the perineural net (PNN), a lattice-like inflexible structure of ECM wrapping certain neurons and dendritic processes (Heo et al., 2018). Taken together, the microscopic and molecular data underscore the ability of exercise to promote remodeling and refinement of the neuronal compartment in iWAT by promoting neuritogenesis.

**Figure 3.**
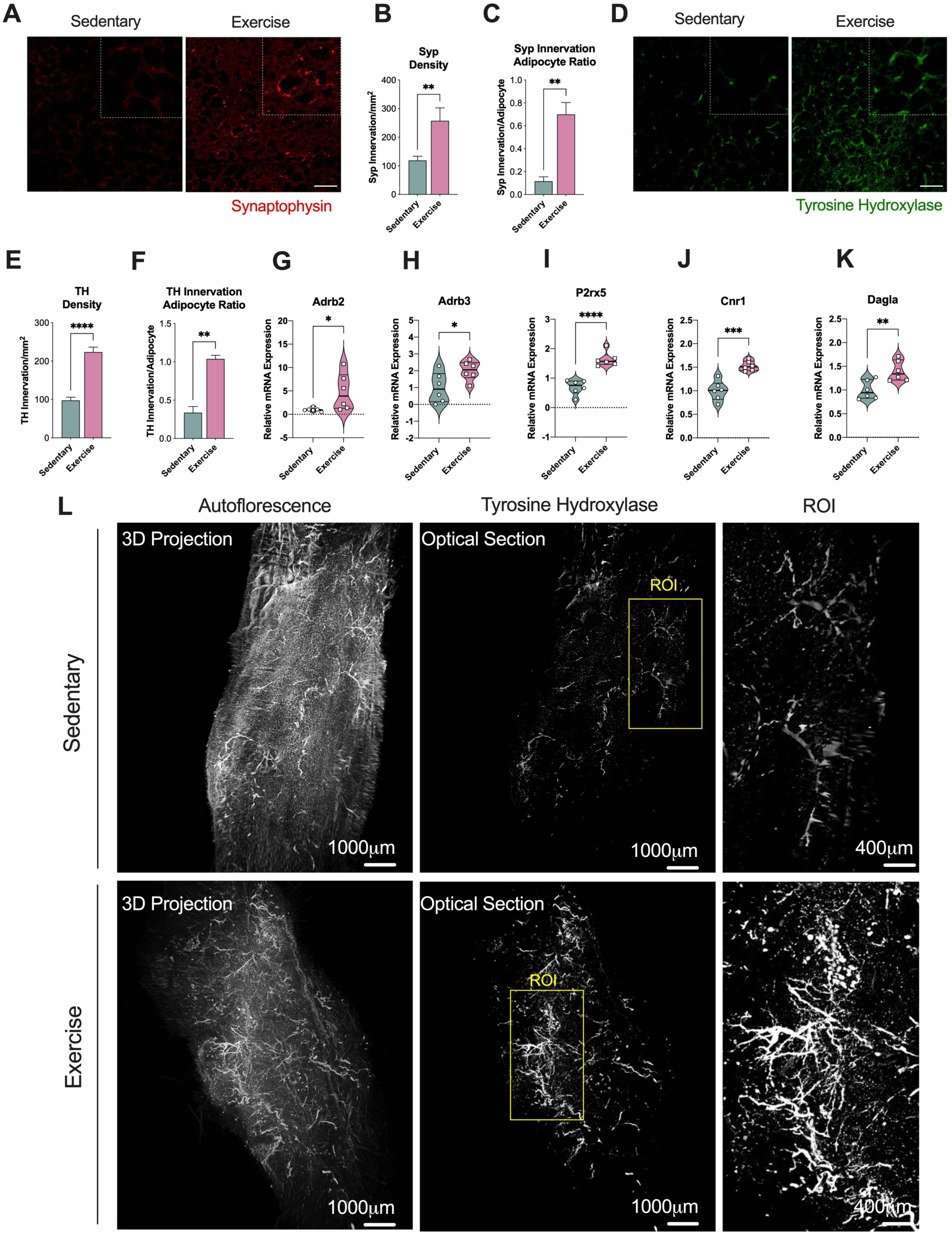
Exercise training increased innervation of iWAT. (A) Representative immunofluorescence staining images of iWAT for the pan-innervation marker Synaptophysin. Scale bar: 50 µm (B-C) Quantification of synaptophysin density (B) and synaptophysin innervation per adipocyte ratio (C) in iWAT from sedentary and trained mice (*n=6/group; calculated from 10 fields/mouse*). (D) Representative immunofluorescence staining images of iWAT for the sympathetic innervation marker Tyrosine Hydroxylase (TH). Scale bar: 50 µm (E-F) Quantification of TH density (E) and TH innervation per adipocyte ratio (F) in iWAT from sedentary and trained mice (*n=6/group; calculated from 10 fields/mouse*). (G-K) Relative mRNA expression of innervation markers *Adrb2* (G), *Adrb3* (H), *P2rx5* (I),*Cnr1* (J), *Dagla* (K) in iWAT from sedentary and trained mice (*n=6/group*). (L) Representative whole-tissue images of iWAT from sedentary (*top*) and trained (*bottom*) mice immunolabeled with TH. Maximum intensity projection from a 1000 μm z-stack and High-magnification view of the region of interest (ROI) are shown. Data are presented as mean ± SEM and were compared using unpaired two-tailed Student’s t test. *p < 0.05, **p < 0.01, ***p < 0.001, and ****p < 0.0001.

### Neuronal Growth Regulator 1 (NEGR1) is an exercise training-induced cell adhesion molecule involved in neuritogenesis of iWAT

To understand the mechanism by which exercise training promotes neuritogenesis we used our omics datasets to identify exercise-regulated proteins known to be involved in neurite outgrowth and synapse formation. We performed a correlation analysis of our proteomics and transcriptomics data and of all the innervation-related proteins, we identified Neuronal Growth Regulator 1 (NEGR1),a member of the IgLON superfamily of cell adhesion molecules (CAMs), as one of the most up-regulated exercise-induced molecules at both the mRNA and protein level (Figure 4A). These findings were validated using qPCR and western blot analysis (Figure S6A-C). We also confirmed the increase of NEGR1 in a separate cohort of male C57BL/6 mice that underwent voluntary wheel training for 10 weeks, where training increased NEGR1 under conditions of both chow (20% fat) and high fat diet (60% fat) (Figure S6D).

**Figure 4.**
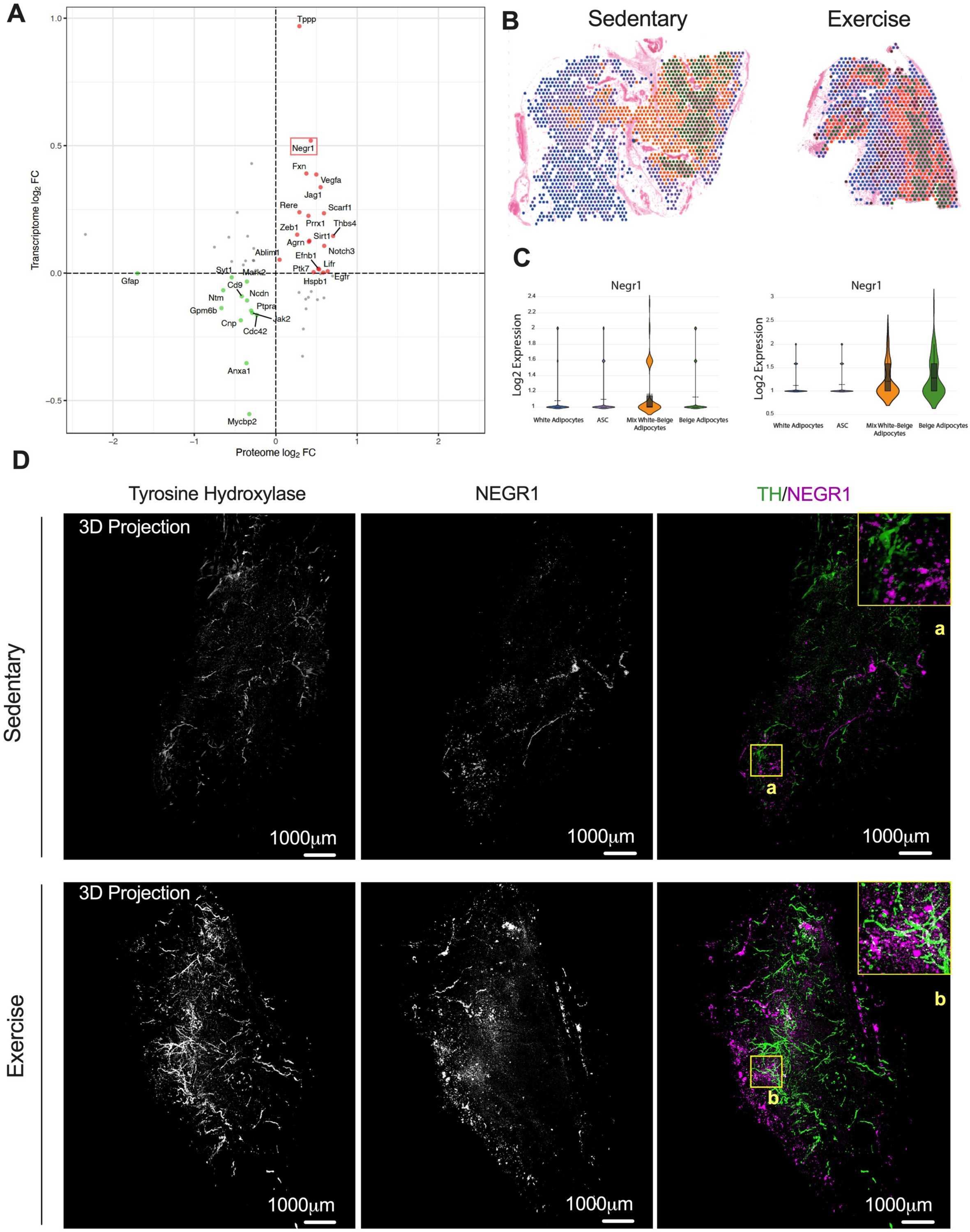
NEGR1 is a cell adhesion molecule induced by exercise training in iWAT. (A) Scatter plot showing correlation of RNA and protein expression in iWAT after 11 days of exercise training. *Negr1* is highlighted with a red box. (B-C) Visium images (B) and relative individual violin plots (C) showing the *Negr1* expression level across the cell clusters detected in iWAT from sedentary (*left*) and exercise training (*right*). (D) Representative whole-tissue images of iWAT from sedentary (*top*) and trained (*bottom*) mice immunolabeled with TH (*green*) and NEGR1(*magenta*). Maximum intensity projection from a 1000 μm z-stack and High-magnification view of ROI are shown (a-b).

We determined the effects of exercise training on the spatial expression of NEGR1 in a cell-type-specific manner to distinguish whether training increased NEGR1 expression in neurons or mature adipocytes. We compared *Negr1* expression in the annotated spatial transcriptomics images (Figure 4B,C) and acquired high-resolution volumetric fluorescent imaging of a full pad of iWAT stained with TH using Adipo-Clear (Figure 4D). Our spatial transcriptomics data demonstrated an increased level of *Negr1* expression in the exercised sample (Figure 4C), with beige adipocytes being the main contributors to *Negr1* expression. These beige adipocytes were neighboring the nerves and vasculature (Figure 4B). To corroborate this finding, we isolated mature adipocytes and ASCs from sedentary and trained iWAT (Figure S6E). Mature adipocytes were immediately homogenized for protein extraction, while ASCs were cultivated until passage 1 (P1) and differentiated into mature adipocytes before mRNA analysis (Figure S6E). We analyzed the mature adipocytes and found an increase of NEGR1 protein levels with exercise training (Figure S6F). ASCs from sedentary and trained mice did not exhibit any differences in *Negr1* mRNA levels (Figure S6G). In contrast, we observed a significant increase of *Negr1* in differentiated adipocytes (Figure S6G). Immunofluorescence in Figure S6H shows expression of NEGR1 in cells with lipid droplets (MERGE) demonstrating that expression occurs primarily in differentiated adipocytes. This figure also shows that Negr1 expression is higher in adipocytes from trained mice compared to sedentary mice. Using 3D projection images, we observed that there was an increase of NEGR1 expression with exercise in the areas with rich sympathetic innervation and this increase was predominantly observed in mature adipocytes and not in neurons (Figure 4D). An animated version of the 3D projection images from the trained mice is shown in Video S1. We pinpointed exercise-induced NEGR1 expression to white/beige adipocyte clusters neighboring the nerves, suggesting that it may function as a neuronal growth regulator at the adipose tissue level.

We then evaluated the effects of exercise training on NEGR1 expression in human subcutaneous WAT (scWAT) collected from two different cohorts before and after supervised exercise training. Study 1 included obese women (n=8, BMI 37.8 +/- 4.7kg/m^2^) who performed 12 weeks of treadmill exercise training (Figure 5A,B). Training significantly improved VO_2_max, and subjects lost ~2kg of body weight without a significant change in % fat mass (Figure 5B). Exercise training increased NEGR1 mRNA expression in both abdominal and gluteal scWAT (Figure 5C,D). Study 2 included lean males (n=10) who performed 12 weeks of combined moderate and high intensity endurance cycling 5 days/week as previously described (Figure 5E) (Yfanti et al., 2010). Analysis of microarray data showed that exercise training significantly increased NEGR1 levels in abdominal scWAT (GSE116801) (Figure 5F) (Takahashi et al., 2019). Results from both these human studies showed an increase in NEGR1 with exercise training regardless of sex, BMI, training period and fat depot. Taken together, based on our mouse and human data, we conclude that NEGR1 is an exercise-induced protein and maybe a key regulator of neuritogenesis in iWAT following exercise training.

**Figure 5.**
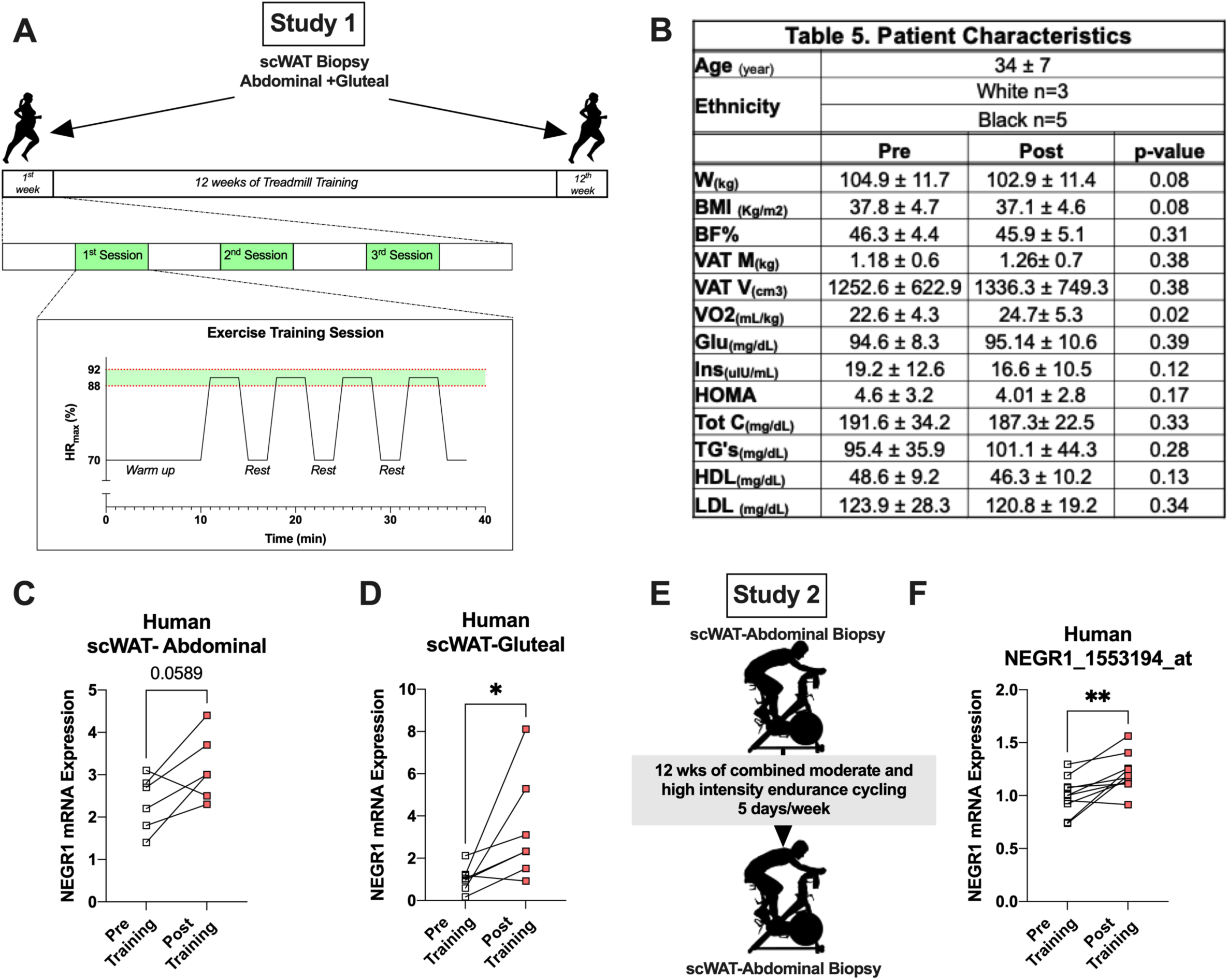
NEGR1 increased with exercise training in human scWAT. (A)Study design to collect subcutaneous WAT (scWAT) biopsies from obese women (*n=6/8*) pre and post treadmill exercise training. (B)Table of patient characteristics enrolled for this study. (C-D)Relative mRNA expression of NEGR1 in abdominal (C) and gluteal (D) scWAT pre and post exercise training (*n=6*). (E) Study design to collect abdominal scWAT biopsies from lean men (*n=10*) pre and post moderate-intensity endurance cycling exercise. (F) Relative mRNA expression of NEGR1 in abdominal scWAT pre and post exercise training (*n=10*) from the microarray dataset GSE116801 Data are presented as mean ± SEM and were compared using paired Student’s test and Two-way ANOVA followed by Tukey’s multiple comparisons test,t, *p < 0.05, and **p < 0.01.

### Adipo-PRDM16KO mice lack exercise-induced iWAT remodeling

To explore the molecular mechanisms involved in iWAT remodeling with exercise training, we investigated Prdm16 because of its critical role in tissue remodeling (Wang et al., 2019). *Prdm16* is a transcriptional regulator that creates a complex with *Gtf2ird1* and *Ehmt1* to repress expression of pro-fibrotic genes (Hasegawa et al., 2018). *Gtf2ird1* is an essential component of this complex and plays a role in inhibiting fibrosis and promoting sympathetic neurite density in scWAT (Hasegawa et al., 2018; Chi et al., 2018b, 2021). We studied both Adipo-cre PRDM16 knockout (PRDM16KO) and wild type mice following 11 days of voluntary wheel running and found no difference in running distance between genotypes (Figure 6A). In wild type mice exercise training did not alter *Prdm16* and *Ehmt1* expression levels but led to a significant increase in *Gtf2ird1* (Figure 6B).

**Figure 6.**
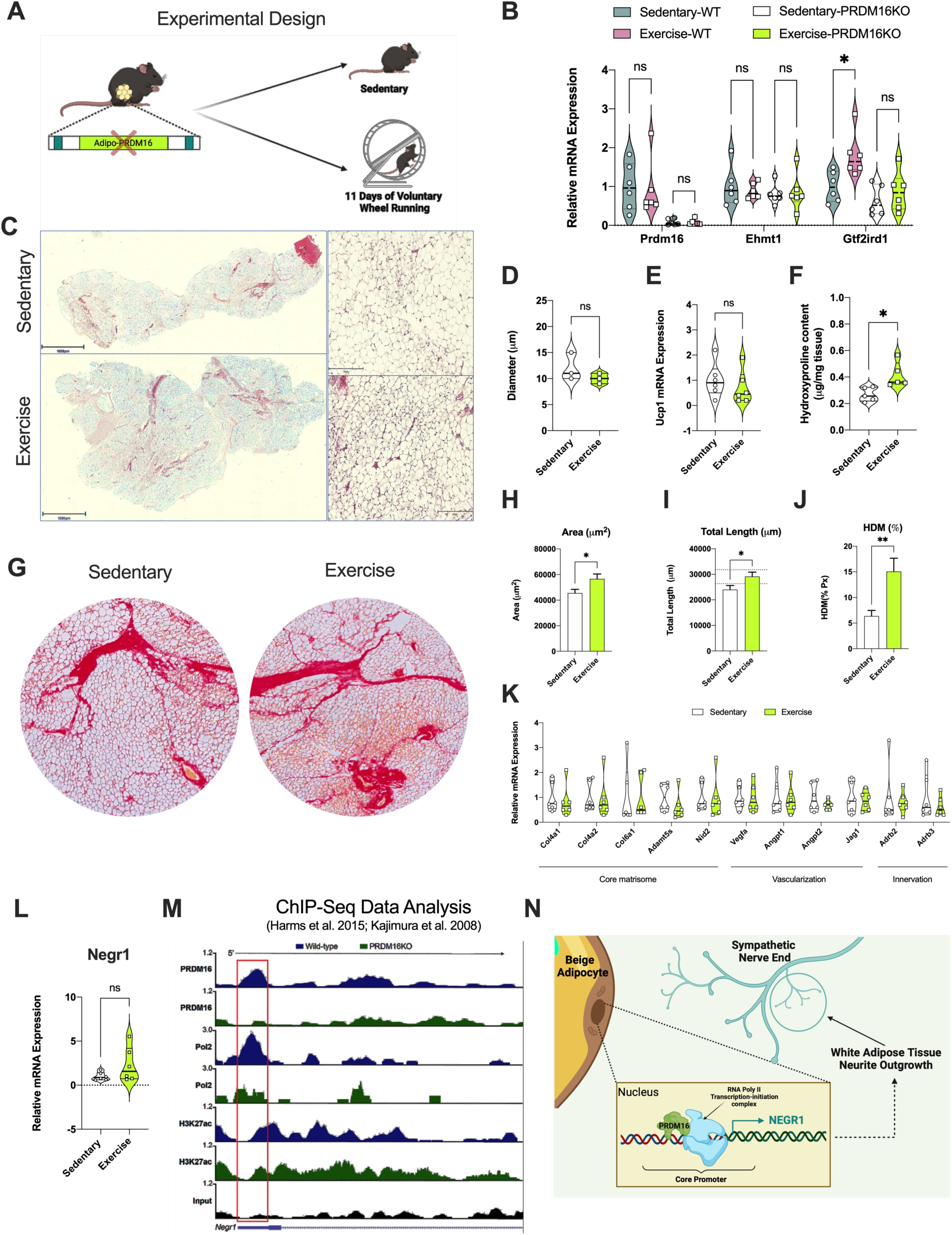
PRDM16 transcriptional complex mediated exercise-induced iWAT remodeling. (A) Study design used for the PRDM16KO mice (*n=6/group*). (B) Relative mRNA expression of the PRDM16 transcriptional complex genes, *Prdm16*, *Ehmt1* and *Gtf2ird1*in iWAT from sedentary and trained Wild Type (WT) and PRDM16KO mice (*n=6/group*). (C-D) Representative H&E staining images (C) of iWAT from sedentary and trained mice with adipocyte cell size measurement (D) (*n=3/group; calculated from 10 fields/mouse*). Scale bar: 500 µm(4x) and 50 µm (20x), respectively. (E) Relative mRNA expression of *Ucp1* in iWAT from sedentary and trained PRDM16KO mice (*n=6/group*) (F) Hydroxyproline content in iWAT from sedentary and trained PRDM16KO mice (*n=5/group*). (G) Representative Sirius Red staining images of iWAT from sedentary (*left*) and trained (*right*) PRDM16KO mice. Scale bar: 100 µm (10x). (H-J) Quantification of ECM deposition pattern area (H),total length of fibers (I), and percentage of high-density matrix (HDM) (J) in iWAT of sedentary and trained PRDM16KO mice (*n=4/group; calculated from 8 fields/mouse*). (K) Relative mRNA expression level for core matrisome, vascularization and innervation markers in iWAT of sedentary and trained PRDM16KO mice (*n=6/group*). (L) Relative mRNA expression of *Negr1* in iWAT from sedentary and trained PRDM16KO mice (*n=6/group*) (M) ChIP-seq profiles in reads per million total reads (RPM) for *PRDM16*, *RNA Polymerase II* (Pol2) and *H3K27ac* in iWAT at *Negr1* for wild type (*blue*) and PRDM16 knockout (*green*) mice. (N) Summary cartoon for the transcriptional regulation of *Negr1* in beige adipocytes and its potential role in the white adipose tissue neurite outgrowth. Data are presented as mean ± SEM and were compared using unpaired two-tailed Student’s t test and Two-way ANOVA followed by Tukey’s multiple comparisons test, *p < 0.05, and **p < 0.01.

We next investigated the phenotype of PRDM16KO and the effects of knocking out *Prdm16* on training-induced ECM remodeling in iWAT. Within the PRDM16KO group, body weight was similar between the trained and sedentary mice, while caloric intake was higher in the trained group (Figure S7A,B). Trained PRDM16KO mice had lower blood glucose levels after 6hrs of fasting compared to sedentary mice (Figure S7C). In the PRDM16KO mice, exercise training did not result in changes to adipocyte size compared to sedentary (Figure 6C,D). Interestingly, lack of PRDM16 prevented an exercise-induced increase in *Ucp1* mRNA levels (Figure 6E) and the presence of multilocular adipocytes, a hallmark of the beiging process. We found increased hydroxyproline content in iWAT of trained PRDM16KO mice compared to the sedentary (Figure 6F). These results contrast the reduced hydroxyproline content found in the trained WT mice (Figure 1C). This finding was confirmed using Sirius Red staining (Figure 6G) and relative quantification analysis in the PRDM16KO iWAT (Figure 6H-J). We found an increase in fibrosis deposition in terms of area, total length and HDM. There were no changes in mRNA expression of core matrisome genes or genes involved in vascularization and/or innervation in trained PRDM16KO iWAT (Figure 6K). Staining with TH showed that exercise training in PRDM16KO mice did not result in increased sympathetic neurite density (Figure S7D), which may be linked to a lack of increased NEGR1 mRNA expression (Figure 6L). Interestingly, in isolated mature adipocytes from PRDM16KO mice we detected lower expression for both NEGR1 mRNA and protein compared to WT (Figure S7E-G). The reduced *Negr1* expression in mature adipocytes from PRDM16KO mice suggests a potential interplay between the NEGR1 and the PRDM16 transcriptional complex in promoting sympathetic neurite density. Using previously published PRDM16 ChIP-seq data in both brown adipose tissue and WAT from WT and PRDM16KO mice (Harms et al., 2015; Kajimura et al., 2008), we identified a PRDM16-binding site located within the promoter region of the *Negr1* gene (Figure 6M). The promoter region also shows an *RNAPol II* peak that is greatly reduced in PRDM16KO, suggesting that PRDM16 binding promotes NEGR1 transcription (Figure 6M). This evidence corroborates our previous spatial transcriptomic data (Fig. 4B) showing that *Negr1* expression was restricted in iWAT to sites neighboring the nerves and beige adipocytes where PRDM16 is crucial for their development (Ohno et al., 2012; Seale et al., 2011). Using the JASPAR database (Castro-Mondragon et al., 2022), we also found three transcription factor binding sites (TFBS) with high scores for Pparg::Rxr heterodimer within the Negr1 promoter region (Figure S7H). This heterodimer is known to be involved in adipogenesis, lipid metabolism and PRDM16 recruitment (Ahmadian et al., 2013; Kajimura, 2015). Taken together, these data demonstrate the pivotal role of PRDM16 transcriptional complex in modulating fibrosis and sympathetic neural refinement of iWAT in the context of endurance physical exercise. Moreover, we show a direct link between PRDM16 and NEGR1 in beige mature adipocytes that may be responsible for the exercise-induced neurite sympathetic density of iWAT (Figure 6N).

## DISCUSSION

Exercise training can reverse many of the detrimental effects of obesity on metabolic health, and adaptations to WAT have been proposed to play a major role in these beneficial effects of exercise. While exercise training has been shown to alter processes such as adipocyte beiging and secretion of metabolically active adipokines (Nigro et al., 2021; Takahashi et al., 2019; Stanford et al., 2015), to our knowledge there had been no investigation of WAT ECM remodeling in response to exercise. This is an important question because recent data suggest the essential role of ECM remodeling in WAT homeostasis. In obesity, WAT is characterized by an extensive expansion of ECM that leads to fibrosis, tissue inflammation and dysfunction (Ruiz-Ojeda et al., 2019). Here, we show that exercise training changes the structural architecture of iWAT and promotes healthy expansion of the tissue by reducing collagen deposition and inducing vascularization and neuritogenesis.

ECM is a significant modulator of cell behavior, function and fate, and structural alterations in ECM have been associated with the development and progression of numerous diseases including cancer and cardiometabolic disease (Sonbol, 2018; Spencer et al., 2011). Thus, elucidating molecular signals that improve ECM function could lead to novel treatments for diseases, including obesity. Our results, showing that exercise training favorably regulates ECM remodeling, thereby facilitating healthy expansion of WAT, provides a physiological model for identification of key molecular adaptations, that in turn could become therapeutic targets. As such, our data underscore the critical role of adipose stem cells (ASCs) in ECM reorganization and remodeling in iWAT. Based on our findings, we hypothesize that in ASCs, TGF-β/SMAD3 signaling functions as a master regulator of collagen species that control ECM structure and function. Interestingly, there is evidence that the TGF-β/Smads signaling pathway is also involved in mesenchymal stem cell commitment and beiging of WAT (Babaei et al., 2018). The prominent role of ASCs in mediating exercise- and obesity-induced adaptations has also emerged through our recent multi-tissue, single-cell analysis of exercise and obesity in rodents (Yang et al., 2022; Rajbhandari et al., 2019).

At the cellular level, we observe that training increases the degree of heterogeneity, changing the proportion of cell types and states, which reflects the “multitasking” role of WAT. We demonstrate changes in both the cellular proportion and the molecular signatures of mature adipocytes with a significant increase in the beige adipocyte population. Another intriguing finding is that exercise training promotes a shift towards the insulin sensitive white adipocytes (LGA) and reduces the presence of the stressed/hypertrophic adipocytes (LSA). We show that there is high collagen gene expression in LSA and their reduction with exercise training directly reflects the reduction in collagen deposition in trained iWAT. The same white adipocyte subpopulations were identified in a study done in visceral WAT in mice fed a high fat diet (Sárvári et al., 2021). This study showed an opposite phenotype from what we see with exercise training, that is, a dramatic decrease of LGA and an increase in LSA subpopulations as well as a significant upregulation of ECM gene programs in LSA (Sárvári et al., 2021). These findings suggest that at least part of the mechanism by which exercise training has beneficial effects on systemic metabolism is by promoting a healthier phenotype in white adipocytes, an effect mediated by repressing the LSA adipocyte subpopulation that plays a prominent role in ECM deposition.

Although exercise-induced vascular and innervation remodeling has been described in skeletal muscle (Laughlin and Roseguini, 2008), whether this occurs in WAT had not been previously studied. Here, we discover that exercise training results in a robust vascular remodeling and neuronal refinement and we hypothesize that this results in the reconfiguration of WAT to allow for both the metabolic and energy balance requirements of physical activity. According to prior studies, the PRDM16 transcriptional complex, consisting of PRDM16, GTF2IRD1 and EHMT1, is one of the main regulators of neuronal refinement and ECM remodeling in WAT (Chi et al., 2018b; Hasegawa et al., 2018). Given its important role in tissue remodeling, we investigated the effects of exercise training on the PRDM16 transcriptional complex. Interestingly, we found that exercise training did not increase *Prdm16* levels but led to a significant increase in *Gtf2ird1,* an important component of the PRDM16 transcriptional complex. Thus, the effects of exercise training on beiging are likely mediated through the interaction of different transcription factors forming a complex, and not through an effect to increase Prdm16 levels. Our finding that training failed to alter beiging and mRNA expression of core matrisome, vascularization and innervation gene markers in iWAT from PRDM16KO mice underscores the important role of this transcriptional complex in mediating the effects of exercise on iWAT.

Studies in the PRDM16KO mice also led us to identify NEGR1 as a potential link between the PRDM16 transcription factor complex and neurite density. NEGR1 is a membrane protein anchored to glycosylphosphatidylinositol that has primarily been studied in the brain, where it mediates neural cell communication and synapse formation (Joo et al., 2019). NEGR1 has been identified in GWAS as a gene implicated in obesity and was shown to increase by ~1.8 fold after 12 days of adipocyte differentiation *in vitro* (Bernhard et al. 2012), but has not been further studied. We found that NEGR1 is significantly increased in both mouse and human WAT in response to exercise training and is highly expressed in mature adipocytes in proximity to nerve endings, and therefore we hypothesize that NEGR1 is a key regulator of exercise-induced WAT innervation. In addition to neurite outgrowth, we propose that the PDRM16-NEGR1 axis is involved in de novo adipogenesis. We base this hypothesis on the finding that the Pparg::Rxr transcription factor heterodimer, known to be involved in adipogenesis, has the highest enrichment score at the *Negr1* promoter region.

Consistent with the mouse models, our human studies reveal that exercise training increases NEGR1 in subcutaneous WAT from male and female subjects. The importance of NEGR1 in metabolic health is also demonstrated by two different GWAS analyses. In one, there was a strong association between NEGR1 common variants and obesity (Thorleifsson et al., 2009; Willer et al., 2009; Speliotes et al., 2010), and in the second, there was a strong association between a NEGR1 locus variant and moderate-to-vigorous physical activity (Klimentidis et al., 2018). All these data point to NEGR1 as a novel, exercise-induced molecule expressed in adipocytes that may play a significant role in regulating metabolic health. Future identification of an agonist that increases NEGR1 may prove to be a new target for the treatment of obesity and metabolic disease. In addition, studies of NEGR1 in other tissues may reveal a broader role for NEGR1 in mediating beneficial effects of exercise on neural and metabolic health.

In conclusion, we have discovered that exercise training is a potent stimulus of iWAT remodeling through fundamental changes in ECM, vascularization, and innervation. We identify robust cell-type specific adaptations in response to exercise training, that in combination with the structural changes, undoubtedly result in a more healthy adipose tissue phenotype. Finally, the extensive single cell, transcriptomic, and proteomic data, in conjunction with detailed imaging and biochemical analyses, provide a resource for identifying novel iWAT factors that can be used to develop therapeutics for the treatment of metabolic disease.

## STAR**⋆**METHODS

- **KEY RESOURCES TABLE**
- **CONTACT FOR REAGENT AND RESOURCE SHARING**
- **EXPERIMENTAL MODEL AND SUBJECT DETAILS**

- Mouse Models
- Human Subcutaneous Abdominal and Gluteal White Adipose Tissues (WAT)
- **METHOD DETAILS**

- Immunocytochemistry and quantification of innervation and vascularization density.
- Measurement of fibrosis and collagen production.
- Whole mount tissue staining and 3D imaging
- Quantitative proteomics of secretome proteins from adipose tissue organ culture.
- Orbitrap-Based LC-MS/MS Analysis
- RNA Isolation and Quantitative Real-Time PCR
- Computational analysis
- Statistical Analysis

### CONTACT FOR REAGENT AND RESOURCE SHARING

Requests for reagents and resources should be directed to the Lead Contact, Laurie J. Goodyear.

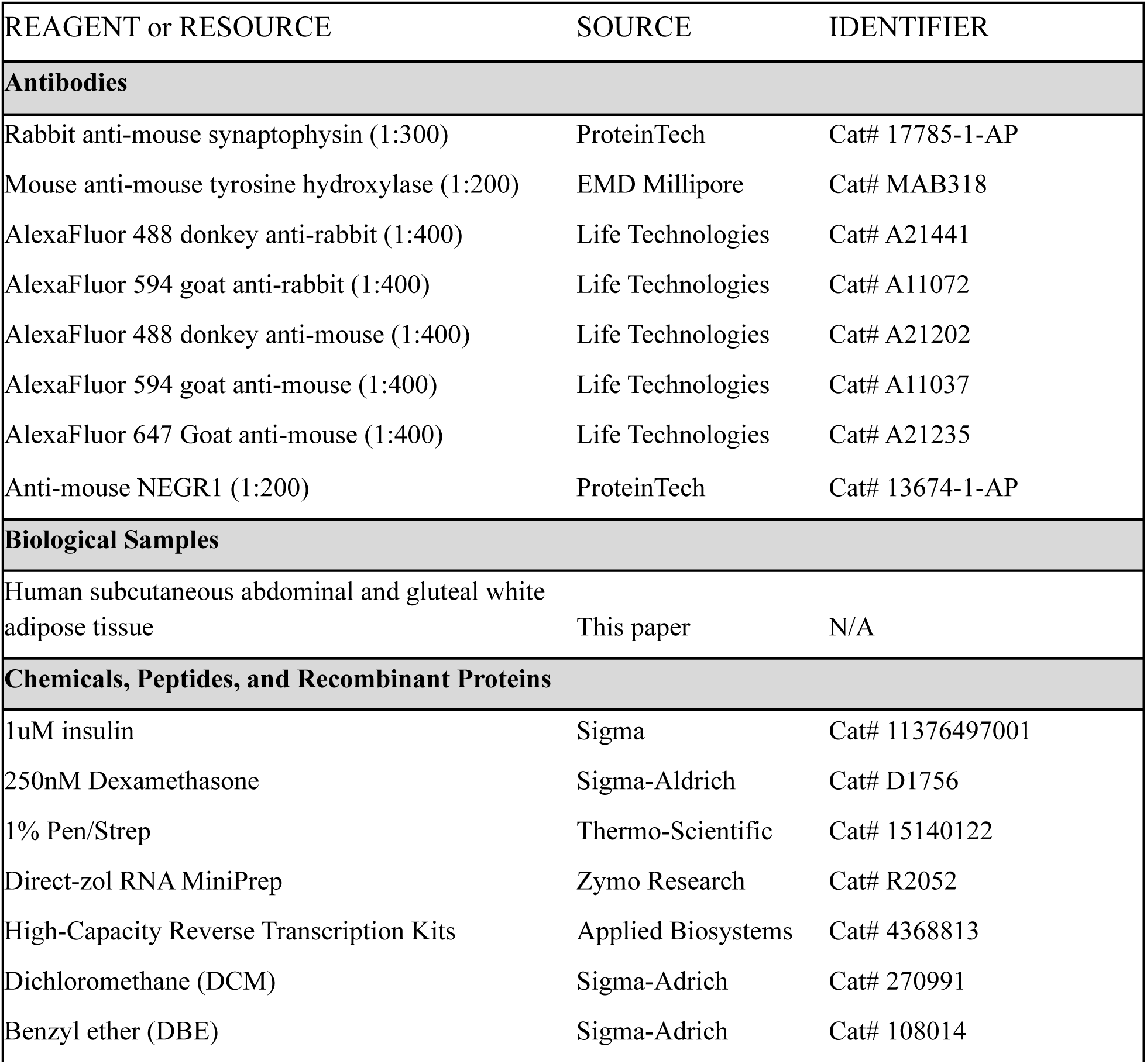
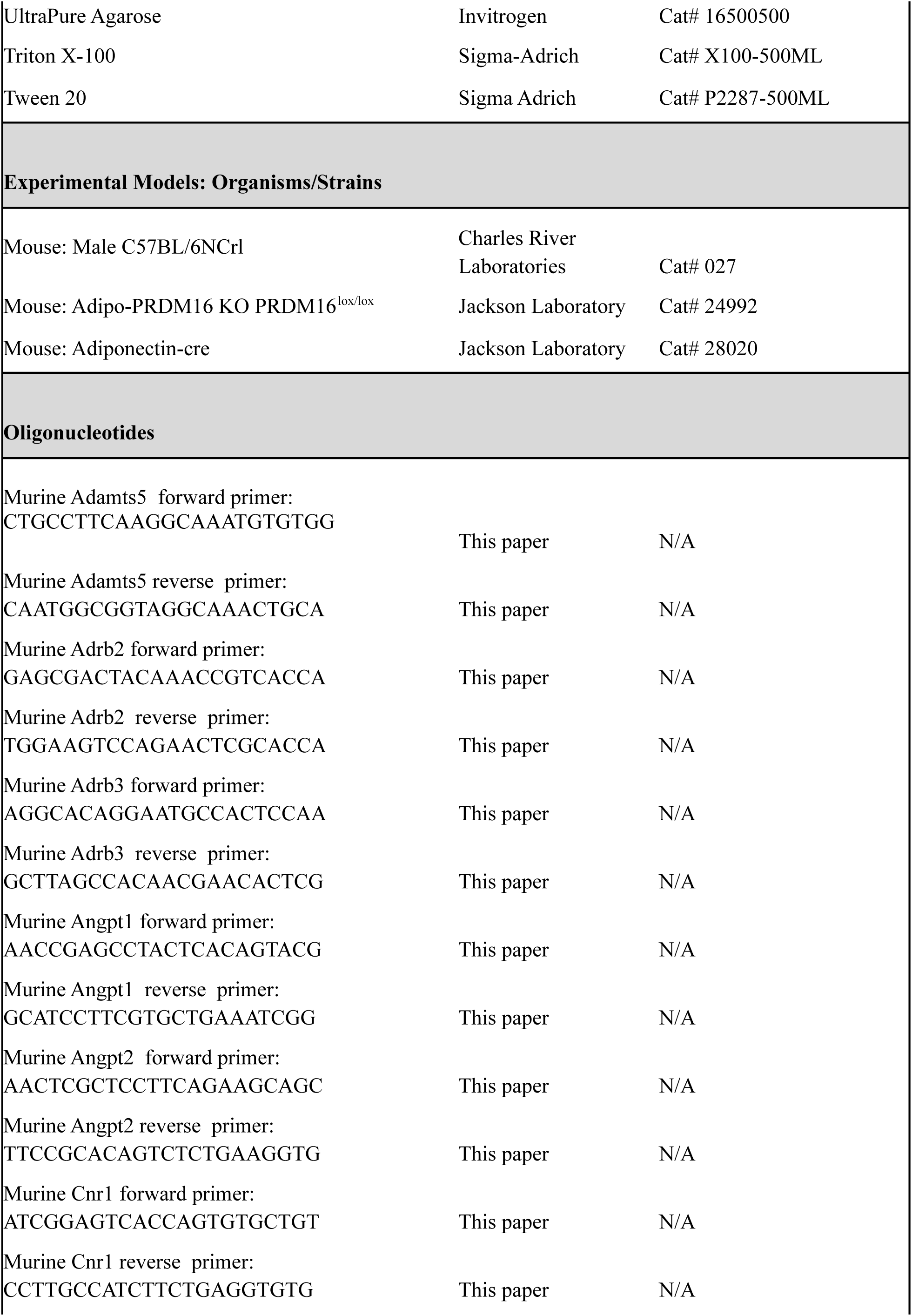
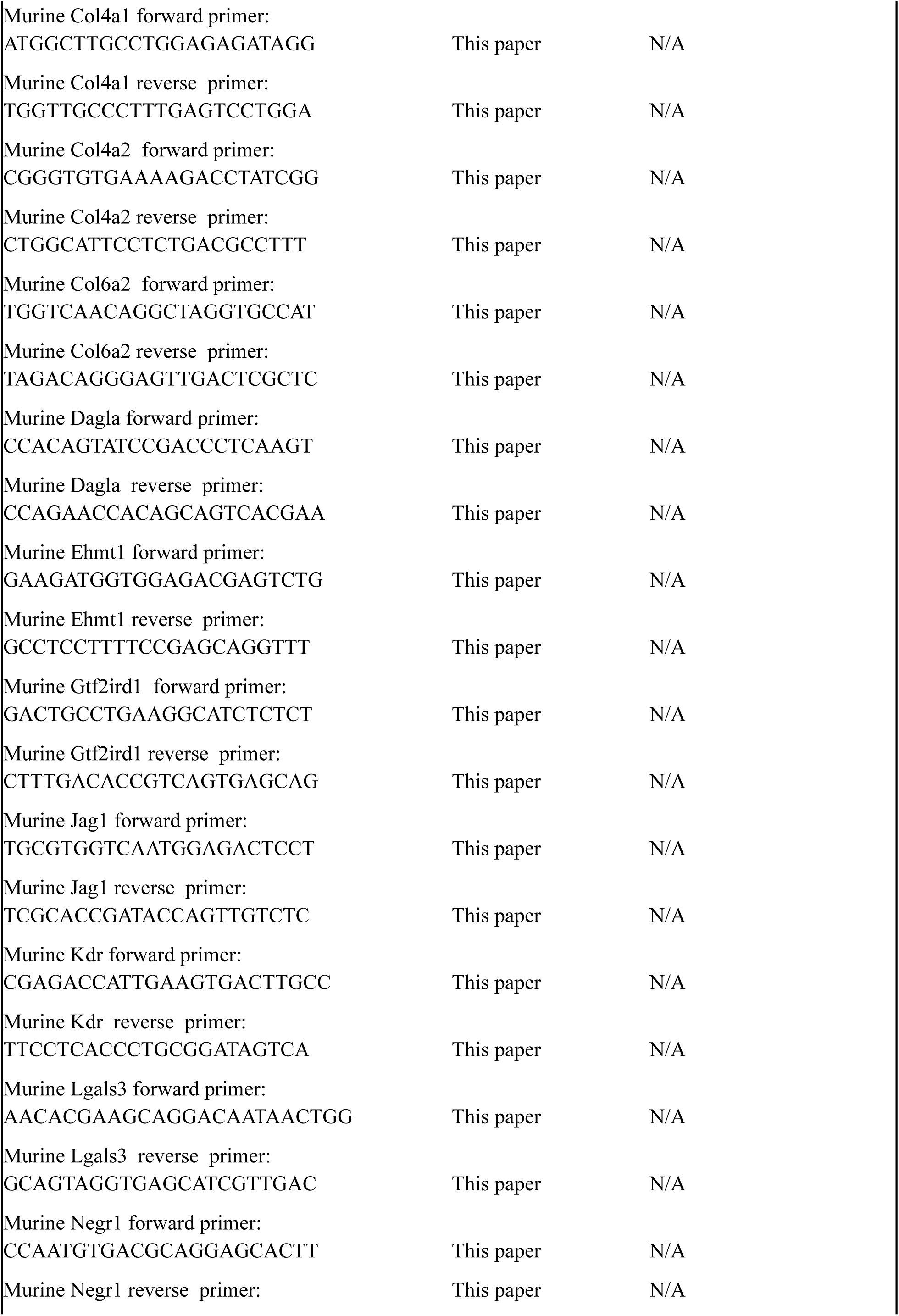
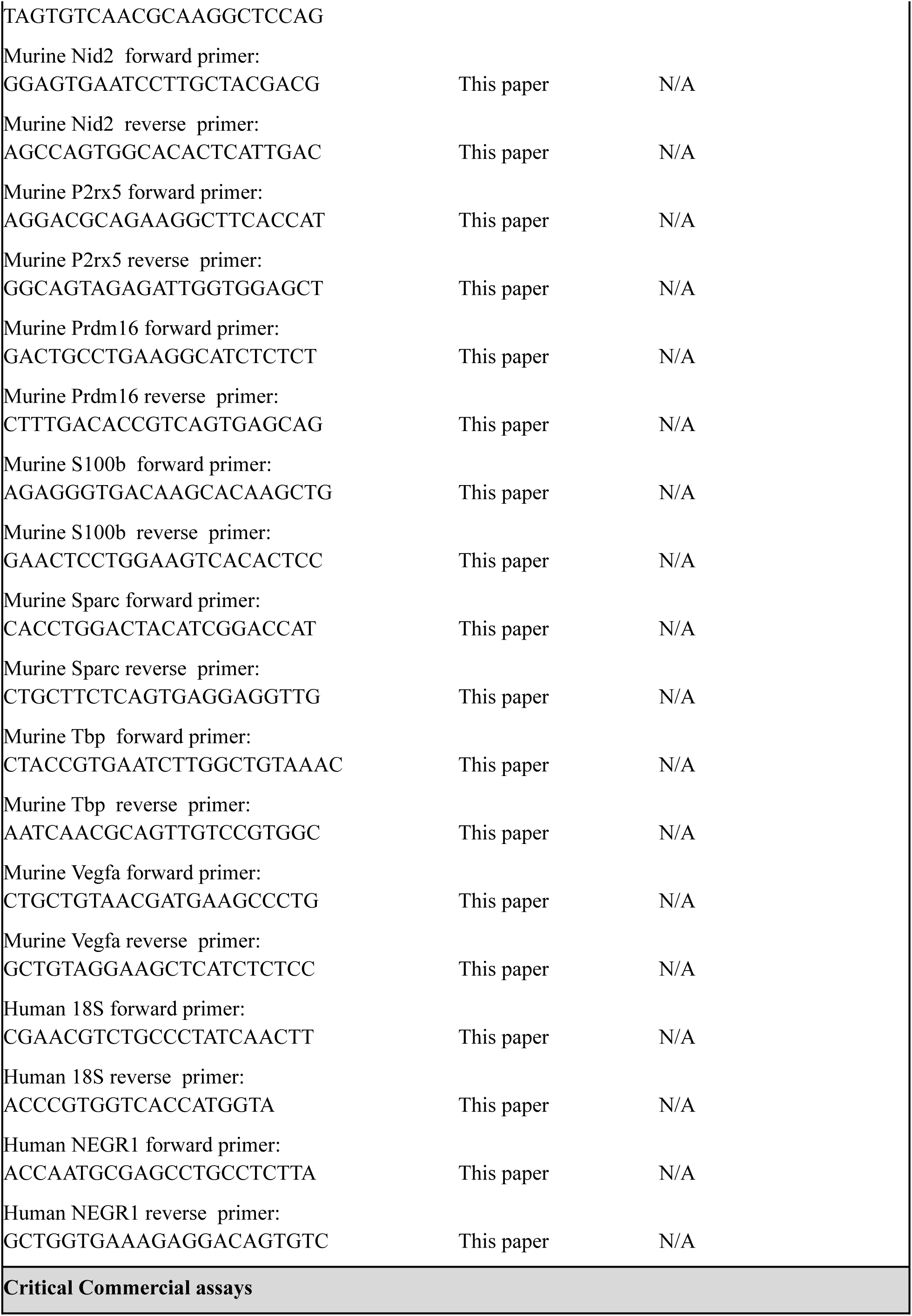
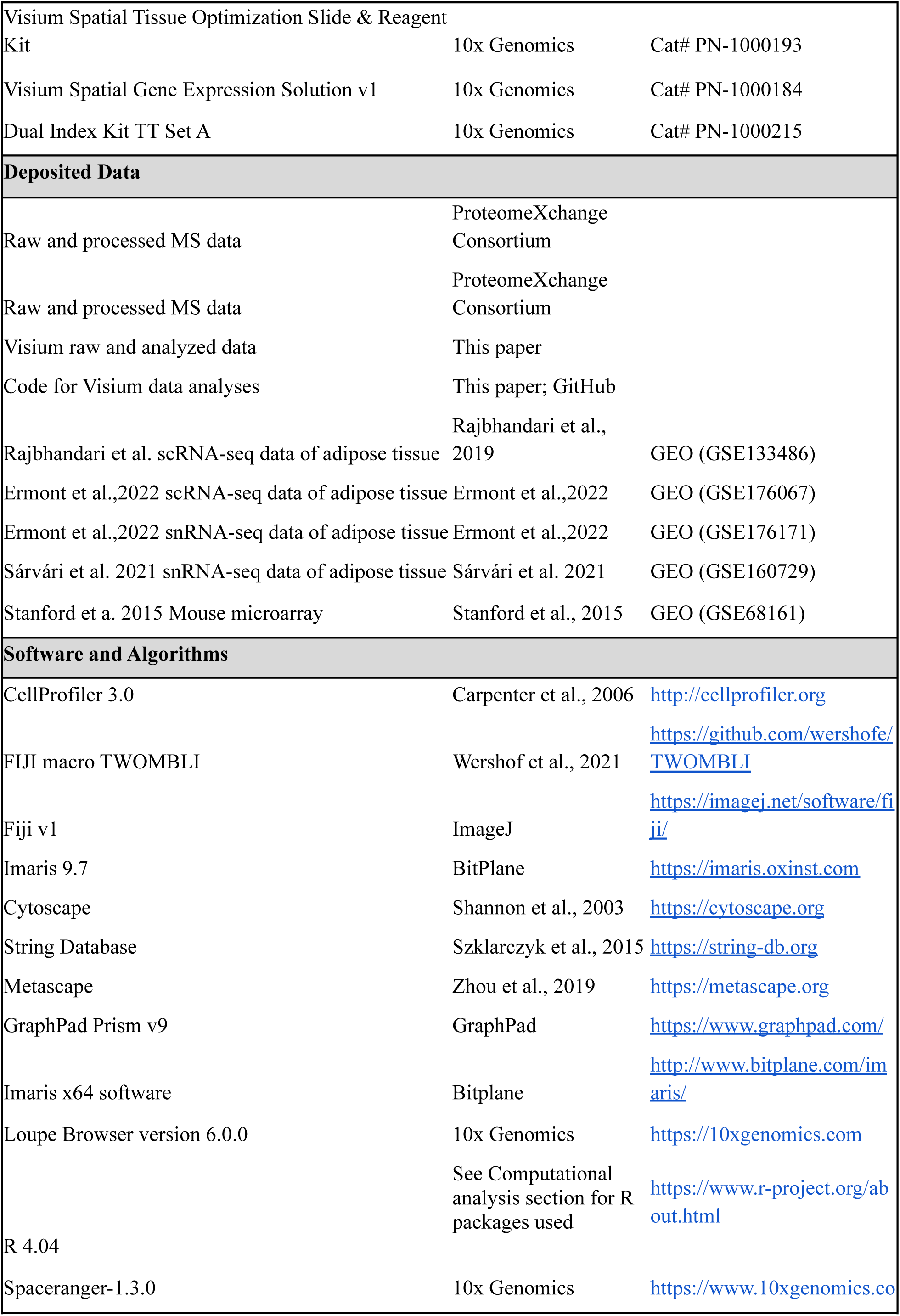
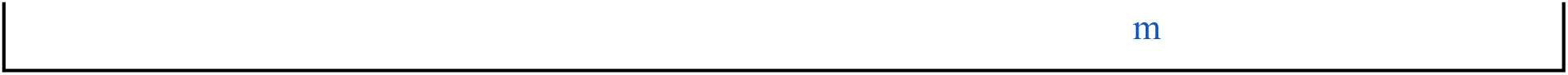

### EXPERIMENTAL MODEL AND SUBJECT DETAILS

#### Mouse Models

All experiments were conducted following NIH guidelines and protocols approved by the IACUC at Joslin Diabetes Center (JDC). 8-week-old male C57BL/6 mice (Charles River Laboratories, MA) were housed at 23°C on a 12/12 h light/dark cycle. Male mice were studied since exercise training in males, but not females, results in beiging adaptations of iWAT. Standard mouse chow diet (9F 5020 Lab Diet, PharmaServ, Inc.) and water were provided ad libitum. For the exercise training experiments, mice were housed with or without a running wheel (24 cm diameter Techniplast) for 11 days. This time period is chosen based on our previous published works showing an optimal training response and maintenance of the adipose tissue depots (Stanford et al. 2015). To generate Adipo-PRDM16 KO PRDM16lox/lox (024992, Jackson Laboratory) mice were crossed with Adiponectin-cre mice (028020, Jackson Laboratory).

#### Human Studies

Both study protocols were approved by the local Ethical Committees: the Institutional Review Board of East Carolina University for Study 1 and Copenhagen and Frederiksberg (KF 01 289434) for Study 2. Both studies were carried out in compliance with the Declaration of Helsinki and received an exempt status by the Institutional Review Board of Joslin Diabetes Center. **Study 1:** Eight women with BMI 37.8 +/- 4.7kg/m^2^ performed a 12-week moderate to high intensity exercise training program (Figure 5A). All subjects had to attend the clinical research center of East Carolina University for three supervised exercise sessions per week using the treadmill. Each exercise session began with a 10 minute warm up followed by four bouts of high intensity interval training of 5 minute duration each during which subjects achieved 88-92% max heart rate. Pre- and post-exercise training subcutaneous WAT biopsies were collected from the abdominal and gluteal area using a biopsy needle. The WAT samples were snap frozen in liquid nitrogen and stored in −80°C until further analysis. **Study 2:** As previously described, a total of 10 young male subjects underwent a 12-week exercise training program which consisted of 60-80 minutes cycling/day. Biopsies of abdominal scWAT were obtained using a biopsy needle before and after the 12 weeks of training (Yfanti et al. 2011). NEGR1 mRNA relative expression was analyzed in human scWAT from the two different studies.

### METHOD DETAILS

#### Immunocytochemistry and quantification of innervation and vascularization density

Inguinal white adipose tissue (iWAT) was fixed in 10% phosphate-buffered formalin and embedded in paraffin. Immunofluorescence was carried out using rabbit anti-mouse synaptophysin (1:300) (17785-1-AP, Proteintech) and mouse anti-mouse tyrosine hydroxylase (1:200) (MAB318, EMD millipore) primary antibodies. Primary antibodies were incubated overnight at 4 °C. The secondary antibodies used were AlexaFluor® 488 donkey anti-rabbit and 594 goat anti-rabbit (A11037, A11072, A11020, A21235, Life Technologies) at a 1:400 dilution, and were incubated for 1 h at room temperature. Images were either captured with an inverted fluorescence microscope (IX51 Olympus) or with a confocal laser scanning microscope (Zeiss LSM 710). Hematoxylin and eosin (H&E) images were acquired with the EVOS M7000 Imaging system (Thermo Fisher System). Synaptophysin, tyrosine hydroxylase and griffonia simplicifolia agglutinin-I were quantified using CellProfiler 3.0 (http://cellprofiler.org). Innervation and vascularization densities were quantified as the number of sprinkles normalized to adipocytes count (mm^2^/adipocytes).

#### Measurement of fibrosis and collagen production

Sirius Red staining was used to quantify the extracellular matrix (ECM) and collagen content in white adipose tissue. Briefly, tissues were fixed overnight in 4% paraformaldehyde and processed for standard paraffin embedding. Sections of 3 μm were stained as previously described (Hemmeryckx et al.) and at least 8 images were captured per slide on an inverted fluorescence microscope (IX51 Olympus) using 10X objective and were analyzed using the FIJI macro TWOMBLI (Wershof et al., 2021). Hydroxyproline content in adipose tissue was measured with a Hydroxyproline Assay Kit (Quickzyme Biosciences).

#### Whole mount tissue staining and 3D imaging

AdipoClear was performed as previously described (Chi et al., 2018a). Anti-mouse TH (1:200) (MAB318,EMD millipore) and anti-mouse NEGR1 (1:200) (13674-1-AP, ProteinTech) were used. Whole-tissue iWAT samples were imaged on a light sheet microscope (Ultramiscroscope II, LaVision Biotec) equipped with 1.6 X objective lenses and an sCMOs camera (Andor Neo). Images were acquired with the Inspector Pro software (LaVision BioTec). Samples were positioned in an imaging chamber filled with dibenzyl ether (DBE) and illuminated from one side by the laser light sheet with 488 and 561 nm laser channels. Whole Tissue 3D images were analyzed using Fiji v1(ImageJ) and Imaris 9.7 (BitPlane).

#### Quantitative proteomics of secretome proteins from adipose tissue organ culture

For the secretome analysis, Quantitative Immunoprecipitation (Quant-IP) analysis was performed. Briefly, 20 mg of iWAT was washed three times in PBS 1X and incubated in the M199 media supplemented with 1µM insulin (11376497001 Sigma), 250nM Dexamethasone (D1756 Sigma-Aldrich) and 1% Pen/Strep (15140122, Thermo-Scientific). After 24hr media was replaced with fresh M199 media plus 1% Pen/Strep. The media was collected after 24hr and immediately stored at −80°C. Debris from the media was removed by centrifugation at 4000 × g. The protein-containing supernatant was mixed with a lysis buffer composed of 50 mM EPPS pH 8, 1 M Urea, 1% SDS, and protease inhibitors (Roche EDTA-free protease inhibitor tablet: 1 mini table per 10 ml). The reduction and alkylation of the tryptic peptides was achieved using 5 mM TCEP, 15 mM Iodoacetamide and 10 mM DTT at 56°C for 30 min. Protein precipitation was done with TCA, precipitate was dissolved 50 mM EPPS. Digestion was performed using Trypsin (1:100) based on protease to protein volume ratio. Each sample was labeled with TMT tags for multiplex proteomics.

#### Orbitrap-Based LC-MS/MS Analysis

Peptides were separated using a gradient of 3 to 36% Buffer B (90% Acetonitrile in 0.1% formic acid) over 120 min. MS analysis was performed using an Orbitrap Fusion^TM^ Tribrid^TM^ Mass Spectrometer (Thermo Scientific, United States) interfaced with an EASY-nLC^TM^ 1200 System. The following 11-plex data search Sequest parameters were used: Peptide Mass Tolerance = 20 ppm, Fragment Ion Tolerance = 1, Max Internal Cleavage Site = 2, Max differential/Sites = 4.

For Reported Quant Parameter: tolerance = 0.003, ms3 = 1, peak picking = max, num isotopes = 2, ms2_isolation_width = 0.7, ms3_isolation_width = 1.2. MS2 spectra were searched using the SEQUEST algorithm against a Uniprot composite database derived from Mouse proteome containing its reversed complement and known contaminants. Peptide spectral matches were filtered to a 1% false discovery rate (FDR) using the target-decoy strategy combined with linear discriminant analysis. The proteins were filtered to a <1% FDR. Proteins were quantified only from peptides with a summed SN threshold of >200 and MS2 isolation specificity of 0.5.

#### Spatial transcriptomics

For spatial transcriptomics we used the 10x Genomics Visium Spatial Gene Expression platform (10x Genomics, Pleasanton, CA, USA). The paraffin-embedded iWAT from sedentary and trained Wild Type and PRDM16KO mice were used for the experiment. Visium Spatial Gene Expression was then carried out according to the manufacturer’s instructions (10x Genomics, Pleasanton, CA, USA) with the following parameters: 20 μm thick sections of adipose tissue were used, H&E staining was carried out by incubating slides for 1 minute in isopropanol, 3 minutes in hematoxylin and 30 seconds in eosin (1:20 dilution). Permeabilization was carried out for 15 minutes. All Visium cDNA libraries were indexed, pooled, and sequenced simultaneously on the Illumina NovaSeq6000 platform, supported by the BioMicro Center Core at MIT. In accordance with the Visium protocol, the number of bases sequenced were 28 nt for R1, 120 nt for R2, and 10 nt for each of the indexes.

#### RNA Isolation and Quantitative Real-Time PCR

RNA was isolated from mouse and human adipose tissue using an RNA extraction kit (Direct-zol™ RNA MiniPrep, Zymo Research). RNA was reverse-transcribed using standard reagents (High Capacity Reverse Transcription Kits, Applied Biosystems), and cDNA was amplified by RT-PCR. For each gene, mRNA expression was calculated relative to Actb and 18s. Primer sequences used in RT-qPCR are presented in Supplementary Table 1.

#### Computational analysis

Proteome and secretome raw data were processed using the pipeline and method to derive fold changes and associated P values for each of the proteins detected as described below. For the tissue secretome, we annotated secreted proteins if the proteins belong to Gene Ontology (GO) terms of either “extracellular region” <u>GO:0005576</u> or “extracellular space” <u>GO:0005615</u>, or the predicted human secretome from <u>Human Protein Atlas</u> by using three methods for signal peptide prediction. We unbiasedly estimated the empirical sample quality weights in the R package limma (Ritchie et al. 2006). Weights varied from 0.4 to 2.5. For both the tissue proteome and secretome, we visualized each sample’s distribution with a boxplot and a histogram to ensure the samples are comparable and that the values are normally distributed. We visualized the clustering of samples in 2D using the first two principal components and performed Surrogate Variables Analysis (SVA) to remove unknown batch effects (i.e. the unwanted variation that is independent of the group assignment) using the R package sva. We performed the principal component analysis (PCA) using the batch-effect-removed data. We assessed if there was a trend between the arrow’s average abundance and its variance. In secretome analysis, we saw a trend and accounted for it in variance estimation using limma-trend (Law et al. 2014). We tested the differential abundance of each row of the expression matrix using limma (Ritchie et al. 2015). We then plotted the proteins from the analysis and if there were multiple comparisons or associations, we selected the proteins that were most significant across these. Volcano and fold-change correlation plots for proteome and secretome analyses were generated using R package ggplot2. The protein-protein interaction analysis was performed using Cytoscape (Shannon et al., 2003) open source software integrating the network analysis generated with String Database (Szklarczyk et al., 2015) and Metascape (Zhou et al., 2019). Spatial data were aligned to the mm10-2020-A reference using spaceranger-1.3.0 with default parameters for each of the samples. We used the R package Seurat to normalize, identify marker genes, reduce dimensions, cluster, annotate cell types and visualize the spatial data. We integrated our spatial data with the publicly available single-cell datasets using FindTransferAnchors and TransferData functions provided by the Seurat package. For spatial distribution analysis we selected region of interests (ROI) in both sedentary and training using Loupe Browser 6. End/PVM cells, Beige, LGA and LSA adipocytes have been identified as cells/dots expressing Cdh5 and Axl log_2_FC > 1, Ucp1 log_2_FC > 4, Pparg and Acaca and Acly log _2_FC > 1, Nnat and Abcd2 log_2_FC > 2 respectively. Control plots have been generated in the same ROI selecting cells expressing Car3 log_2_FC>5. Interaction potential was calculated using the MosaicIA plugin in ImageJ as interaction strength (*ε)* using parametric Hernquist potential (Law et al. 2014; Ritchie et al. 2006; Ritchie et al. 2015)

### Statistical Analysis

Data are expressed as mean ± SEM. Sample sizes are indicated in the figure legends. All statistical analyses were performed using GraphPad Prism v9. Statistical significance was analyzed by One-way ANOVA and two-way ANOVA followed by Tukey’s multiple comparisons test or unpaired two-tailed Student’s t-test.

## ACKNOWLEDGMENTS

This work was supported by NIH grant R01DK099511 and R01DK101043 (to L.J.G.), K23DK114550 (to R.J.W.M), 5T32DK00726042, 1F32DK12643201 and Joslin DRC Pilot and Feasibility Program award (to M.V.), and the Joslin Diabetes Center DRC (P30 DK36836). The Centre for Physical Activity Research (CFAS) is supported by TrygFonden (grants ID 101390, ID 20045, and ID 125132). During the study period, the Centre of Inflammation and Metabolism (CIM) was supported by a grant from the Danish National Research Foundation (DNRF55). We thank Dr Krithika Ramachandran and Madison M. Columbus for comments and scientific feedback. Dr Michael R Blanchard and Dr Mahmoud EL-Rifai for technical assistance with Adipo-Clear images acquisition and analysis. Dr Hui Pan, Dr Jonathan M. Dreyfuss for statistical analysis support.

## AUTHOR CONTRIBUTIONS

P.N. and M.V. designed research, carried out experiments, analyzed data and wrote the paper. J.H. R.C. and N.C. performed genotyping of PRDM16 knockout mice and cell experiments. L.H. coordinated and supervised the spatial transcriptomics analysis. J.Y. T.C and D.P. performed bioinformatics analysis. M.F.H. supervised all experiments. J.D.W., J.R., R.C.H., B.K.P and S.N. carried out and provided human samples. R.J.M. and M.K. analyzed data, wrote the paper. L.J.G. directed the research project, designed experiments and wrote the paper. All authors have participated in the manuscript review. All authors approved the final manuscript.

## SUPPLEMENTARY FIGURE LEGENDS

**Figure S1 Systemic phenotypic effects of 11 days of exercise training on C57BL/6 male mice, with qPCR data for ECM genes, related to** **Figure 1**.

(A-C) Total food consumption (A), body weight change (B), and glucose (C) for all sedentary and trained mice used in the study (*n=12/group*).

(D-E) Serum insulin level (D) and insulin resistance index HOMA-IR (E) for a sub cohort of n=6/group mice.

(F) Overview of exercise training effect on mouse matrisome gene categories in iWAT using microarray dataset (GSE68161).

(G) List of top 10 matrisome associated genes changing with exercise training versus sedentary mice (p<0.05) divided in subcategories: ECM regulators (*green*), ECM affiliated (*yellow*) and secreted factors (*red*).

(H) List of core matrisome genes changing with training (p<0.05) divided in subcategories: glycoproteins (*magenta*), collagens (*pink*), and proteoglycans (*purple*).

(I) Protein-protein interaction (PPI) networks analysis for iWAT matrisome genes changed with exercise training.

(J-K) mRNA expression level for matrisome associated genes (J) and core matrisome genes (K)(*n=6/group*).

(L) Categorization of identified proteins in proteome modulated by exercise training using GO Cellular Component terms.

(M) Pathway analysis for the exercise-regulated proteins using the Matrisome gene sets available in the Molecular Signature Database (MSigDB v5.0).

(N) Categorization of identified proteins in secretome modulated by exercise training using GO Cellular Component terms.

(O) Pathway analysis for the exercise-regulated proteins using the Matrisome gene sets.

(P) Protein-protein interaction (PPI) networks analysis for iWAT conditioned media secreted proteins that are decreased with exercise training.

Data are presented as mean ± SEM and were compared using unpaired two-tailed Student’s t test. *p < 0.05, **p < 0.01, and ***p < 0.001.

**Figure S2 Unsupervised analysis of two independent single-cell transcriptomics datasets, related to** **Figure 2**.

(A) Unsupervised clustering of 85,258 cells from the iWAT of pooled male C57BL/6 mice. 10 distinct cell groups were represented on a UMAP plot.

(B) Individual violin plots showing the expression levels and distribution of representative genes for the core matrisome proteins decreased with exercise training detected in iWAT conditioned media (secretome analysis). The y-axis is the log-scale normalized read count.

(C) Unsupervised clustering of 61,285 cells from the iWAT of pooled male C57BL/6 mice. 13 distinct cell groups were represented on a UMAP plot.

(D) Individual violin plots showing the expression levels and distribution of representative genes for the core matrisome proteins decreased with exercise training detected in iWAT conditioned media (secretome analysis). The y-axis is the log-scale normalized read count.

(E) Inferred TF activity based on target gene expression levels across cell types identified in the single-cell data presented in panel (A). The TFs were ranked by their activity levels in ASCs. The percentage of ECM genes in the targets of the individual TFs was plotted to the left of the heatmap.

**Figure S3 Spatial transcriptomics analysis of iWAT from sedentary and exercise training mice, related to** **Figure 2**.

(A) Violin plots showing the variance in molecular counts across spots and the total number of spots counted in iWAT from sedentary (*left*) and exercise training (*right*) mice.

(B-C) Dotplot visualization listing cell clusters identified with spatial transcriptomics on iWAT from sedentary (B) and exercise training (C) mice. Cell clusters listed on the y-axis, showing unbiased gene expression for the top genes per cluster identified by log fold change; genes (features) are listed along the x-axis. Dot size reflects percentage of cells in a cluster expressing each gene; dot color reflects expression level (as indicated in the legend).

(D) Individual violin plots showing the expression level for basement membrane collagens species across 5 selected cell type clusters.

Data are presented as mean ± SEM and were compared using One-way ANOVA. *p < 0.05, **p < 0.01, ***p < 0.001, and ****p < 0.0001.

**Figure S4 Spatial transcriptomics analysis of iWAT from sedentary and exercise training mice, related to** **Figure 2**.

(A) Probability density distribution plots showing the distance relative to the beige adipocyte from preadipocytes (pre-adipo) in iWAT from sedentary (*left*) and exercise training (*right*) mice. Dashed black lines indicate the random distribution among the clusters.

(B) Interaction analysis for LGA, LSA and Endo/PVM clusters in iWAT from sedentary (*top*) and exercise training (*bottom*). The plots show the observed nearest-neighbor (NN) distance distribution (*blue*) between two arbitrary point patterns, the model fits (*green*) to the observed NN distance distribution and the random context distribution (*red*).

**Figure S5 ECM proteins are key components for vasculature and innervation remodeling in iWAT, related to Figure 4 and 5**.

(A-B) Venn diagram illustrating quantified proteins regulated by exercise training in proteome (A) and secretome (B) datasets, annotated as members of innervation, vascularization and ECM pathways.

(C) Representative images of iWAT from sedentary and trained mice stained with Griffonia Simplicifolia Lectin (GSA-I). Scale bar: 50 µm

(D-E) Quantification of capillary density (D) and capillary per adipocyte ratio (E) in iWAT from sedentary and trained mice (*n=6/group; calculated from 10 fields/mouse*).

(F-J) Relative mRNA expression of pro-angiogenic markers *Vegfa* (F), *Kdr/Vegfr* (G), *Jag1*(H), *Angpt1*(I),*Angpt2*(J) in iWAT from sedentary and trained mice (*n=6/group*).

(K-L) Protein-protein interaction (PPI) networks analysis for iWAT proteins up-regulated (K) and down-regulated (L) with exercise training detected in proteome dataset and annotated in the innervation pathway.

Data are presented as mean ± SEM and were compared using unpaired two-tailed Student’s t test. *p < 0.05, **p < 0.01, and ****p < 0.0001.

**Figure S6 NEGR1 is expressed in mature adipocytes.**

(A) Relative mRNA expression of *Negr1* in iWAT from sedentary and trained mice (*n=6/group*).

(B-C) Representative images of NEGR1 protein by western blot (B) in sedentary and trained mice with relative quantification (C) (*n=6/group*).

(D) Relative mRNA expression of *Negr1* in iWAT from sedentary and trained mice under standard diet (20% fat) and high fat diet (60% fat) (*n=6/group*).

(E) Study design to collect fresh mature adipocytes and adipose stem cells (ASCs) from sedentary and trained mice (*n=12/group, 4 mice were pooled to reach n=3*).

(F) Representative images of NEGR1 protein by western blot in fresh mature adipocytes isolated from sedentary and trained mice with relative quantification (*n=3/group*).

(G) Relative mRNA expression of *Negr1*in undifferentiated ASCs and differentiated mature adipocytes isolated from sedentary and trained mice (*n=3/group*).

(H) Representative images of differentiated mature adipocytes isolated from sedentary and trained mice immunostained for NEGR1, DAPI (*nuclei*) and lipidTOX for neutral lipid stain (*n=3/group*).Scale bar: 50 µm (20x).

Data are presented as mean ± SEM and were compared unpaired two-tailed Student’s t test and Two-way ANOVA followed by Tukey’s multiple comparisons test, *p < 0.05, ***p < 0.001.

**Figure S7 PRDM16 transcriptional complex mediated exercise-induced iWAT remodeling, related to** **Figure 6**.

(A-C) Body weight (A), total food consumption (B), and glucose (C) for sedentary and trained PRDM16KO mice (*n=6/group*)

(D) Representative immunofluorescence staining images of iWAT for the sympathetic innervation marker Tyrosine Hydroxylase (TH). Scale bar: 50 µm

(E) Relative mRNA expression of *Negr1* in fresh mature adipocytes isolated from wild type and PRDM16KO mice (*n=12/group, 4 mice were pooled to reach n=3*).

(F-G) Representative images of NEGR1 protein by western blot (F) in fresh mature adipocytes isolated from wild type and PRDM16KO mice with relative quantification (G) (*n=12/group, 4 mice were pooled to reach n=3*).

(H) The sequence analysis of murine *Negr1* gene promoter region (−2000bp relative to the transcription start site, TSS). The highlighted regions in blue, green and yellow depict the three high scored consensus sequences for the Pparg::Rxra transcription factor complex. Data are presented as mean ± SEM and were compared using unpaired two-tailed Student’s t test, *p < 0.05, **p < 0.01, ***p < 0.001, and ****p < 0.0001.

